# Myeloid HIF-1α Sustains Hypoxic Fibrotic Fronts and Drives Pulmonary Fibrosis

**DOI:** 10.64898/2026.03.09.710578

**Authors:** Yuanyi Wei, Christopher Bailey, Peng Zhang, Jose Herazo-Maya, Theodoros Karampitsakos, Joseph Pescrille, Felisa Diaz-Mendez, Daniel Lee, Andriy Samokhin, Yan Liu, Wonder Drake, Yin Wang

**Affiliations:** Division of Immunotherapy, Institute of Human Virology, University of Maryland School of Medicine, Baltimore, MD 21201, USA; Department of Surgery, University of Maryland School of Medicine, Baltimore, MD 21201, USA; Department of Neurosurgery, Beijing Children’s Hospital, Capital Medical University, National Cancer for Children’s Health, Beijing, China; Division of Pulmonary, Critical Care and Sleep Medicine, Department of Medicine, University of South Florida, Morsani College of Medicine, Tampa, FL 33612; Department of Medicine University of Maryland School of Medicine, Baltimore, MD 21201, USA; Department of Microbiology and Immunology, University of Maryland School of Medicine, Baltimore, MD 21201, USA

**Keywords:** Pulmonary fibrosis, Hypoxia, HIF-1α, Macrophage-fibroblast interactions, Lipid nanoparticle, Echinomycin, Fibrogenesis

## Abstract

**Rationale:** Progressive fibrosing interstitial lung disease features “advancing fronts” where new matrix is deposited, but the signals sustaining these propagating niches remain incompletely defined.

**Objectives:** To determine the spatial and temporal compartments in which HIF-1α operates during fibrotic progression and to test whether myeloid HIF-1α is a tractable driver of lesion propagation.

**Methods:** We integrated human IPF datasets, clinical severity profiling, sarcoidosis peripheral blood immune phenotyping, multiplex immunofluorescence and spatial mapping in human lung tissue, single-cell transcriptomic analyses, and temporally staged bleomycin lung injury with genetic and lung-directed therapeutic perturbations.

**Measurements and Main Results:** In the Lung Genome Research Consortium cohort, HIF1A expression was increased in IPF lungs and correlated with higher GAP scores. In sarcoidosis, circulating monocytes from patients with progressive disease exhibited increased HIF-1α compared with those with resolving disease. In IPF lungs, nuclear HIF-1α localized predominantly to CD68⁺ macrophages and PDGFRα⁺ fibroblasts concentrated within collagen-rich, αSMA⁺ advancing fronts, and single-cell analyses demonstrated enrichment of HIF-1α-linked transcriptional programs consistent with macrophage-fibroblast crosstalk (including pro-fibrotic growth factors, chemokines, and matrix-regulatory pathways). In bleomycin-induced fibrosis, HIF-1α activity emerged first in macrophages and subsequently in fibroblasts within pimonidazole-marked hypoxic rims bordering nascent αSMA⁺ foci. Myeloid-specific Hif1a deletion reduced front-associated macrophage persistence, attenuated fibroblast activation, and decreased collagen deposition. Two lung-directed strategies, inhaled liposomal echinomycin and inhaled shHif1a lipid nanoparticles, phenocopied these effects, demonstrating therapeutic tractability.

**Conclusions:** These findings define a hypoxic front-zone niche in which myeloid HIF-1α sustains macrophage persistence and promotes fibroblast activation and matrix remodeling. By linking spatial compartmentalization to causal genetics and lung-directed intervention, our work identifies myeloid HIF-1α as a mechanism-anchored, locally targetable driver of fibrotic lesion propagation.

## Introduction

Progressive interstitial lung disease (ILD) is characterized by relentless extracellular matrix accumulation, distortion of alveolar architecture, and irreversible loss of lung function (1, 2). Current antifibrotics, pirfenidone and nintedanib, slow physiologic decline but rarely halt progression, underscoring the need for mechanism-anchored and spatially precise therapeutic strategies (2-4).

Histopathology and spatial profiling support a front-core organization of usual interstitial pneumonia in which advancing fibrotic fronts at lesion edges are major sites of collagen biosynthesis, whereas central regions are more consolidated and relatively acellular (5, 6). This suggests that programs operating at fibrotic fronts may be most relevant to lesion propagation and that mapping cell states and signals within this niche may reveal actionable targets.

Across organs, fibrosis is increasingly viewed as a spatiotemporally organized process in which macrophage programs evolve over time and localize to discrete tissue niches, thereby coordinating fibroblast activation and matrix remodeling (7). Hypoxia is a defining feature of fibrotic tissue and stabilizes hypoxia-inducible factors (HIFs), which regulate metabolic and transcriptional adaptation across multiple cell types (8-10). In macrophages, HIF-1α promotes glycolytic rewiring and effector programs that shape persistence and cytokine output, while in mesenchymal compartments HIF signaling is linked to matrix production and altered collagen structure-function (8-10). However, the compartment and timing in which HIF-1α acts to sustain lung fibrosis, and the most tractable node for intervention, remain insufficiently defined.

To test whether myeloid HIF-1α maintains a hypoxic, fibroblast-activating niche at advancing lesion fronts, we integrated human IPF tissue and dataset analyses with temporally staged bleomycin injury (11) and conditional myeloid genetics using LysM-Cre (12). We complemented these approaches with lung-directed pharmacologic and RNA-interference perturbations of the HIF-1α pathway to assess therapeutic tractability (13, 14), with hypoxia verified by pimonidazole labeling.

In this study, we test the hypothesis that myeloid HIF-1α sustains a hypoxic niche that supports fibroblast activation at advancing fibrotic fronts. By integrating human IPF profiling with temporally staged injury and complementary lung-directed modes of HIF-1α perturbation, we define a spatially and temporally organized front-zone program of myeloid HIF-1α and assess whether selectively disrupting this axis linking hypoxia, macrophages, and fibroblasts can limit fibrotic progression in vivo.

## Results

### HIF-1α is elevated in IPF lung, localizes to macrophage and fibroblast niches, and correlates with disease severity

We previously reported that increased programmed death signaling on immune cells is associated with poor prognosis in sarcoidosis and IPF (15). To further evaluate the clinical relevance of HIF1A in interstitial lung disease, we analyzed HIF1A expression in IPF lungs from the LGRC cohort. HIF1A was significantly higher in IPF (n=123) than controls (n=96) [10.69 (95% CI: 10.59-10.81) vs 9.96 (95% CI: 9.87-10.04), p<0.0001] (Fig. 1A) and positively correlated with GAP score (n=111, r=0.3120, 95% CI: 0.13-0.47, p=0.0009) (Fig. 1B,C). Cohort demographics are shown in Table S1. Immunohistochemistry confirmed increased HIF-1α protein in fibrotic regions, with prominent nuclear staining in lesion-associated cells (Fig. 1D). Multiplex immunofluorescence identified CD68⁺ macrophages as a major HIF-1α⁺ cellular source and demonstrated co-localization with profibrotic mediators (TGFβ, VEGF) and Collagen I-rich regions (Fig. 1E). In sarcoidosis lung, IL-6 signal co-localized with CD68⁺ macrophages (Fig. S1A) and was frequently juxtaposed with HIF-1α⁺ cells (Fig. S1B). Consistent with a myeloid-biased hypoxia program, PBMC flow cytometry from sarcoidosis patients showed higher intracellular HIF-1α in monocytes than in T or B cells (Fig. S1C).

**Fig. 1.**
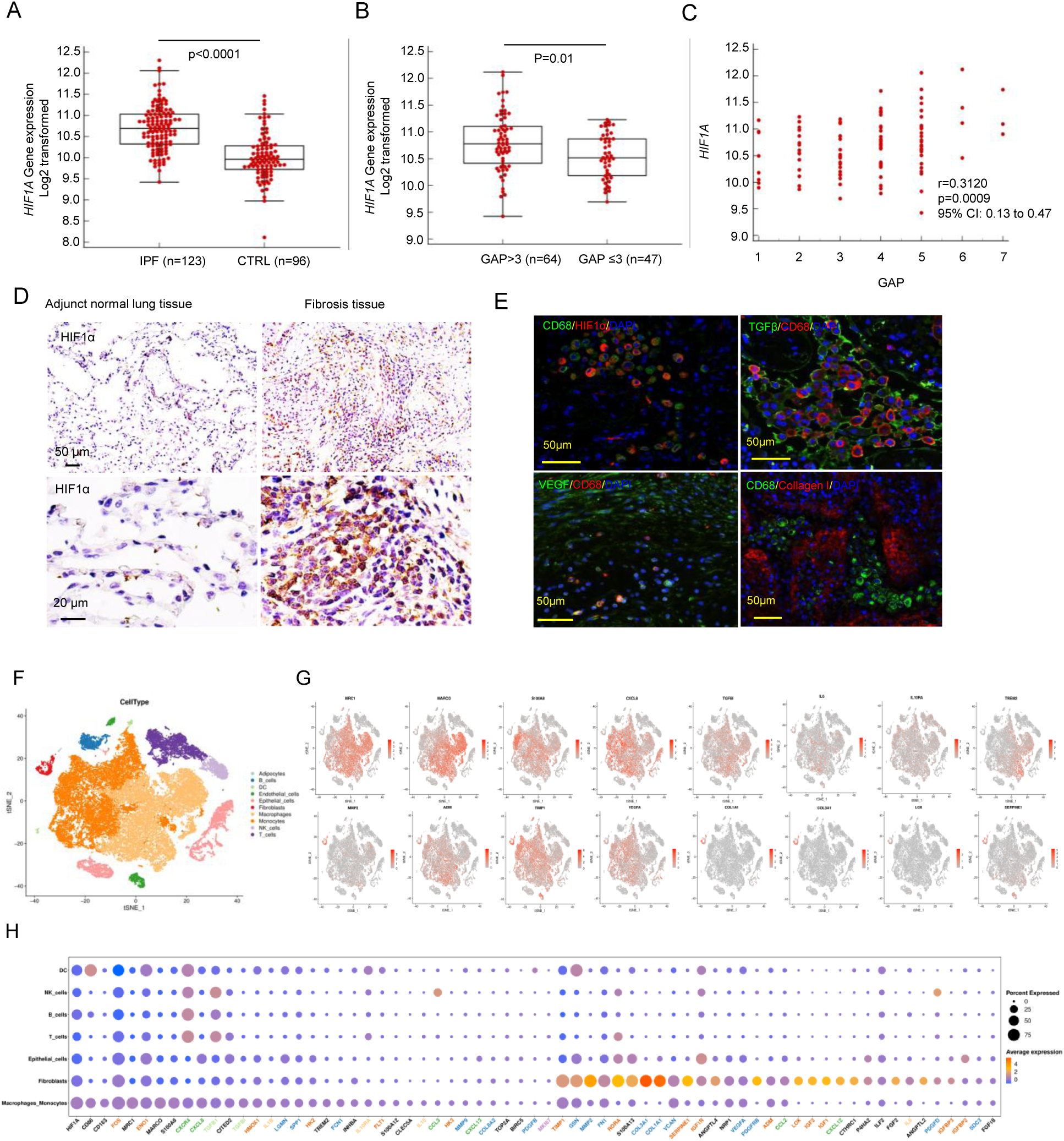
Macrophages/monocytes and fibroblasts adopt a hypoxic response signature in fibrotic tissues. **A.** LGRC dataset analysis showing significantly higher *HIF1A* expression in IPF lungs (n=123) compared with controls (n=96). **B.** Subgroup analysis of IPF patients stratified by GAP stage demonstrates higher HIF1A expression in patients with advanced disease (GAP >3; n=64) compared with those with lower GAP scores (GAP ≤3; n=47). **C.** Correlation between HIF1A expression and GAP score in IPF lungs (n=111), showing a positive association. **D.** Immunohistochemical staining for HIF-1α in human IPF lung tissue versus adjacent normal lung tissue. IPF lungs show robust nuclear HIF-1α staining in fibrotic regions. **E.** Immunofluorescence co-staining of IPF lung sections showing CD68^+^ macrophages colocalized with HIF-1α, TGFβ, VEGF, and Collagen I, indicating activation and pro-fibrotic mediator production. **F.** tSNE (t-Distributed Stochastic Neighbor Embedding) / UMAP (Uniform Manifold Approximation and Projection) representation of 35,500 single-cell transcriptomes in 32 IPF lungs and 28 control donor lungs. Cell-type annotation was based on expression of canonical marker genes. **G.** Marker gene expression and mRNA counts, color-coded and projected onto the tSNE embedding in (F). **H.** Dot plot of scaled, log-normalized expression of marker genes of the clusters. Gene names color-coded by functional categories. Dot size indicates percentage of cells per cluster with any mRNAs detected, and color shows Z-scores of log-normalized mRNA counts.

To define HIF-1α-linked programs across lung cell types, we analyzed scRNA-seq data (GEO GSE136831) (16) from 32 IPF and 28 donor lungs. HIF-1α target gene enrichment was highest in monocyte/macrophage and fibroblast compartments (Fig. 1F), which comprised dominant fractions of the analyzed cells (Fig. 1F). Within these populations, monocytes/macrophages expressed profibrotic mediators and inflammatory factors including TGFB1, PDGF, VEGFA, and IL1B, while fibroblasts expressed extracellular matrix and remodeling-associated genes (Fig. 1G,H) (17, 18). Ligand–receptor analysis highlighted growth factor and cytokine signaling pathways (e.g., TGFB1-TGFBR1/2, PDGF-PDGFR, VEGFA-VEGFR, IL1B-IL1R1) and chemokine axes implicated in macrophage–fibroblast communication and recruitment (Fig. 1G,H) (17, 18). These programs include multiple known HIF-1α-regulated genes in macrophages and fibroblasts that support angiogenesis, inflammation, and matrix remodeling (19-23). Together, human tissue and single-cell analyses place HIF-1α activity within macrophage- and fibroblast-rich niches linked to profibrotic signaling and disease severity.

### Temporal Ordering of Macrophages, Fibroblasts, and HIF-1α in Bleomycin Lung Fibrosis

To define the temporal appearance of macrophages, fibroblasts, and HIF-1α signaling during bleomycin-induced fibrosis, we analyzed single-cell lung suspensions by flow cytometry across harmonized time bins: Day 1 (baseline), Day 3 (early inflammation/early fibrogenesis), Days 5-7 (active fibrogenesis), and Days 10-14 (fibrosis) (Fig. 2A-D). HIF-1α signaling can promote fibrogenic remodeling via mediators such as PDGF and VEGF (24, 25). Monocyte-derived alveolar macrophages (MoAMs) increased by Day 3 and remained elevated through Days 10-14 (Fig. 2A). MoAMs were defined using a published gating strategy as SiglecF^int^ CD11c^+^ CD64^+^ macrophages, distinct from AM (SiglecF^hi^ CD11c^hi^) and IM (SiglecF^−^ CD11c^−/low^) (26). In contrast, PDGFRα⁺ fibroblast subsets were low at Day 3 and increased from Days 5-7 through Days 10-14 (Fig. 2B-C), including both PDGFRα⁺CD26⁺CD90⁻ and PDGFRα⁺CD26⁻CD90⁻ populations, indicating that fibroblast expansion lags macrophage accumulation by approximately 2-4 days. We also quantified lymphocytes and neutrophils. CD3⁺ T cells and CD19⁺ B cells were significantly higher at Days 10-14 than Days 1-3 (Fig. S2A-D), with CD4⁺ and CD8⁺ T cells showing the same late expansion pattern (Fig. S2B,C). Neutrophils increased by Days 5-7 and remained elevated at Days 10-14 (Fig. S2E).

**Fig. 2.**
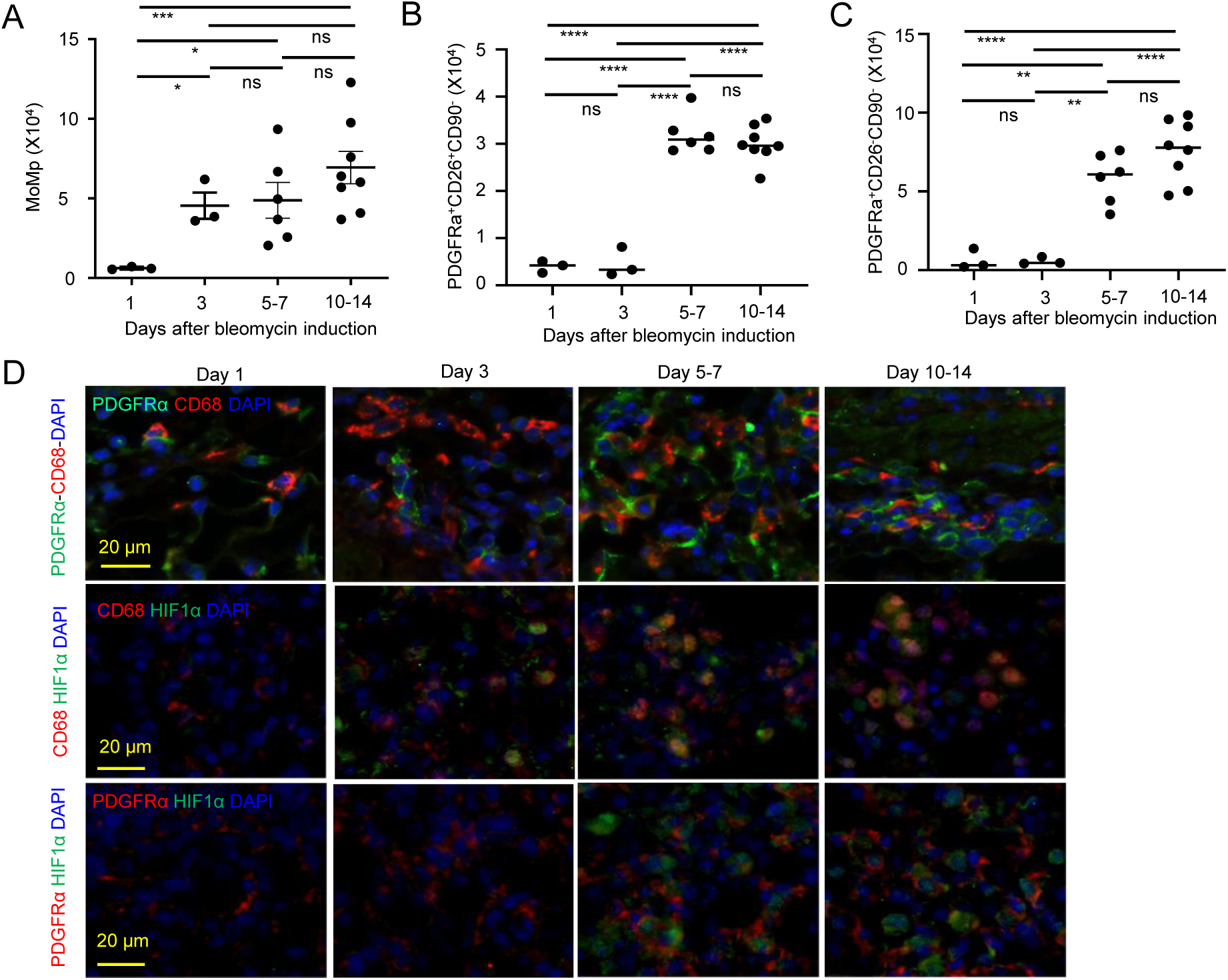
Temporal dynamics of macrophage and fibroblast subsets in the lung following bleomycin-induced pulmonary fibrosis. Flow cytometric quantification of myeloid and fibroblast populations at indicated days after bleomycin administration. **A-C.** Quantification across harmonized bins: Day 1 (Baseline), Day 3 (Early inflammation and fibrogenesis), Days 5-7 (Active fibrogenesis), and Days 10-14 (Fibrosis). **A.** Monocyte-derived alveolar macrophages (MoAMs) increase at Day 3 and remain elevated at Mid and Late. **B-C.** PDGFRα⁺ fibroblast subsets rise later. (B) PDGFRα⁺CD26⁺CD90⁻ (activated/pro-matrix) and (C) PDGFRα⁺CD26⁺CD90⁺ (differentiated/myofibroblast-leaning) increase beginning Days 5–7 and remain high Days 10-14. Points are individual mice; bars show mean ± SEM. p < 0.05 (*), < 0.01 (**), < 0.001 (***), < 0.0001 (****); ns, not significant. **D.** Representative multiplex IF across time bins. Top: PDGFRα with CD68 and DAPI; middle: CD68 with HIF-1α; bottom: PDGFRα with HIF-1α. HIF-1α staining is detectable in CD68⁺ cells by Day 3 and becomes evident in PDGFRα⁺ fibroblasts from Days 5-7 onward. Data are representative of two independent experiments.

Multiplex immunofluorescence revealed CD68⁺/HIF-1α⁺ cells by Day 3, coincident with the MoAM increase (Fig. 2D). By Days 5-7, PDGFRα⁺/HIF-1α⁺ fibroblasts were evident and persisted through Days 10-14 (Fig. 2D), paralleling the flow-cytometric expansion of PDGFRα⁺ subsets. Thus, across matched bins, macrophage accumulation occurred first (Day 3), fibroblast expansion followed (Days 5-7), and HIF-1α staining emerged within each compartment during its respective rise. This temporal ordering is descriptive and motivates subsequent functional tests of HIF-1α pathway disruption *in vivo*.

### Hypoxic macrophages concentrate at fibrotic fronts while mature scar cores are macrophage-limited

Serial immunofluorescence in bleomycin-injured mouse lung demonstrated a reproducible spatial coupling of hypoxia, macrophages, and myofibroblasts at nascent fibrotic foci. Pimonidazole staining (Probe) outlined hypoxic rims encircling small αSMA⁺ islands (Fig. 3A, insets I-IV), indicating a hypoxic border around emerging myofibroblast clusters. In an adjacent serial section, many hypoxic Probe⁺ cells co-localized with CD68 (Fig. 3B), and CD68 staining confirmed a perilesional macrophage cuff surrounding the same αSMA⁺ structures (Fig. 3C), positioning macrophages at the advancing fibrotic front. This spatial pattern places hypoxic/HIF-1α^+^ macrophages at the lesion edge in close proximity to emerging αSMA^+^ myofibroblast foci (27-29). These imaging data do not by themselves establish causality, but they associate front-zone hypoxic macrophages with areas of fibroblast remodeling (28, 29).

**Fig. 3.**
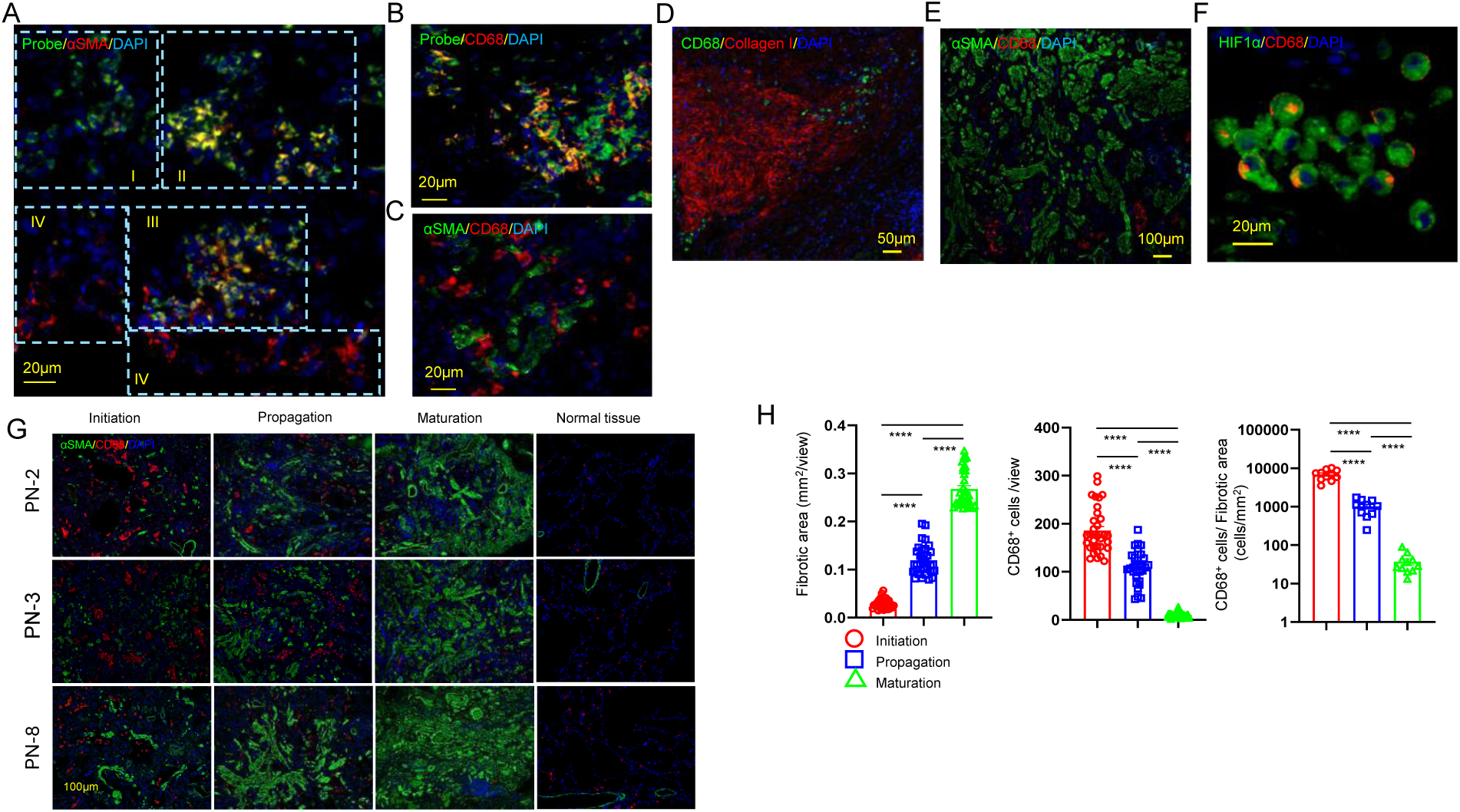
Macrophages are hypoxic and concentrate at the fibrotic front in mouse and human lung fibrosis. **A-C.** Mouse lung sections at different fibrotic phases after bleomycin injury. Hypoxyprobe (pimonidazole) was administered 90 minutes before euthanasia on day 14 post-BLM induction to assess hypoxic regions. **A.** Multiplex immunofluorescence showing αSMA⁺ myofibroblasts (red) and hypoxia probe signal (green) with DAPI nuclear staining (blue). Distinct fibrotic phases (I-IV) are indicated. **B.** Consecutive section of (A) stained for CD68⁺ macrophages (red) demonstrates their enrichment within hypoxic niches co-localized with hypoxyprobe signal (green). **C.** Consecutive section of (A) showing αSMA⁺ myofibroblasts (green) in close spatial association with CD68⁺ macrophages (red). **D-H.** Human IPF lung tissue. **D.** CD68⁺ macrophages (green) overlap with collagen I deposition (red) in fibrotic regions. **E.** αSMA⁺ myofibroblasts (green) with interspersed CD68⁺ macrophages (red) in advanced fibrosis. **F.** High-magnification view showing CD68⁺ macrophages (red) co-expressing HIF-1α (green). **G.** Multiplex IF across three patients (PN-2, PN-3, PN-8) staged as Initiation (small αSMA foci with macrophage “swarming”), Propagation (expanding plaques with macrophage cuffing), Maturation (established αSMA cores with sparse intralesional CD68), and Normal tissue. **H.** Quantification across staged lesions. Left: Fibrotic area (mm²/view) increases from Initiation to Propagation to Maturation. Middle: CD68⁺ cells per view are highest in Initiation, intermediate in Propagation, and lowest in Maturation. Right: CD68⁺ cell density per fibrotic area (cells/mm²; log scale) shows the same decreasing trend across stages. Bars show mean ± SEM; **** P < 0.0001. Data are representative of three independent experiments.

To test whether this spatial organization extends to human disease, we examined IPF lungs. Collagen-dense cores contained few intralesional macrophages, whereas CD68⁺ cells persisted at lesion margins and along septa (Fig. 3D), with wide-field αSMA/CD68 imaging showed macrophages outlining αSMA-dense regions (Fig. 3E). HIF-1α signal was prominent in CD68⁺ macrophages within IPF tissue (Fig. 3F). Clinical metadata for the IPF cohort used in Fig. 3E are summarized in Table S2. To assess generalizability across cases, we examined additional fields from all nine patients (Fig. S2). Across cases, αSMA/CD68 staining reproduced the same spatial pattern observed in Fig. 3E: CD68⁺ macrophages concentrated at lesion margins and interstitial interfaces, with reduced intralesional signal in mature, collagen-dense cores. This case-level survey confirms that macrophage enrichment at the fibrotic front and relative depletion within consolidated scar are consistent features across the cohort. We then staged human lesions into Initiation (small αSMA⁺ foci with macrophage “swarming”), Propagation (expanding plaques with macrophage cuffing at the edge), and Maturation (consolidated αSMA⁺ cores with sparse intralesional macrophages), with normal tissue as reference (Fig. 3G; Table S2). Fibrotic area increased stepwise across stages, whereas CD68⁺ cells per view and CD68⁺ cell density normalized to fibrotic area decreased (Fig. 3H). In sarcoidosis lungs, CD68⁺ macrophages localized to ECM-rich remodeled regions and a subset was HIF-1α⁺ (Fig. S4), supporting generalizability across fibrotic lung contexts. Together, the mouse and human datasets support a phase-specific redistribution in which macrophages, including a hypoxia-associated HIF-1α-positive fraction, are enriched at nascent and expanding αSMA⁺ fronts while mature scar cores are relatively macrophage-poor. This pattern aligns with prior reports localizing profibrotic macrophage populations to remodeling regions in pulmonary fibrosis (27-29), and with spatial characterizations of IPF lesion evolution (6, 30).

### *Hif1a* Deletion Reduces Macrophage Accumulation and Attenuates Bleomycin-Induced Pulmonary Fibrosis

Given the dual role of macrophages as both drivers and regulators of fibrotic lung disease, there is a current emphasis on exploring therapeutic strategies targeting pro-fibrotic macrophages in both pre-clinical and clinical studies (31). To assess the contribution of HIF-1α in myeloid cells during fibrogenesis, we used mice with conditional *Hif1a* deletion in myeloid lineages driven by LysM-Cre (32). Remarkably, flow cytometry showed a reduction in the percentage of monocyte-derived macrophages (MoMp) in BAL after bleomycin, with no significant change in alveolar macrophages (Fig. 4A), using a previously described gating strategy (26). In digested lung tissue, both MoMp and alveolar macrophages were reduced in *Hif1a* knockout mice compared with wild-type (WT) controls (Fig. 4B). These data indicate that myeloid *Hif1a* loss is associated with fewer pulmonary macrophages during bleomycin-induced fibrogenesis. Immunofluorescence for CD68 in fixed lung specimens further confirmeded fewer CD68⁺ cells at the interface between normal parenchyma and fibrotic foci in *Hif1a* knockout mice than in WT (Fig. 4C-D), consistent with decreased macrophage accumulation at the fibrotic front. Consistent with macrophage-fibroblast crosstalk in fibrosis, this reduction is expected to blunt fibroblast activation and matrix deposition, aligning with the decreased αSMA⁺ remodeling observed in *Hif1a*-deficient lungs (Fig. 4E). In contrast, neutrophil numbers in BAL and lung tissue did not differ significantly between genotypes (Fig. S5), suggesting a relatively selective effect on macrophage-lineage cells.

**Fig. 4.**
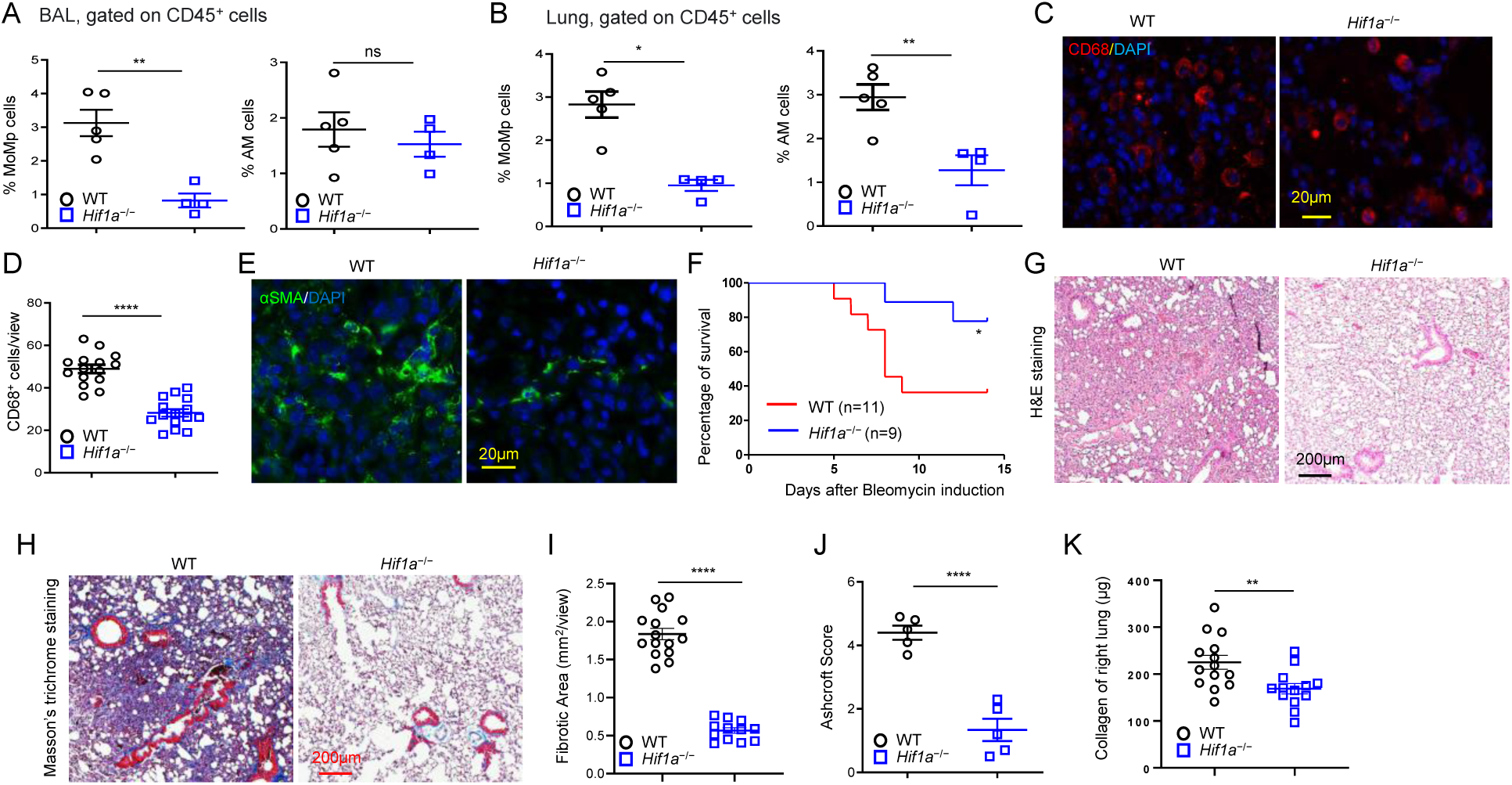
Targeted mutation of the *Hif1a* gene reduces macrophages in BAL and lung and mitigates bleomycin-induced pulmonary fibrosis in BLM-induced mice. **A.** Flow cytometry analysis of BAL samples from WT or *Hif1a*^−/−^ macrophages after BLM administration. The percentage of WT and *Hif1a*^−/−^ macrophages gated on CD45^+^ subsets in BAL at 14 days post-BLM administration, shown as mean ± SEM. Data are representative of 3 independent experiments. **B.** Frequency of macrophages gated on CD45^+^ subsets in digested lung tissues analyzed through flow cytometry from WT and *Hif1a*^−/−^ BLM-induced mice. MoMp: CD45^+^CD64^+^MHC-II^-^CD11b^+^; AM: CD45^+^SiglecF^+^CD11c^+^CD64^+^. The summarized data are presented as mean ± SEM for one experiment and are representative of 3 independent experiments. **C, D.** Immunofluorescence staining with anti-mouse CD68 reveals a decreased number of macrophages in the lung from *Hif1a*^−/−^ BLM-induced mice compared with the WT group (C). Macrophage counts in the lung were quantified at 40× magnification based on CD68 immunofluorescent staining, with n = 3 fields per lung from each of 5 mice (D). **E.** Immunofluorescence staining with anti-αSMA antibody to examine α-SMA^+^ myofibroblasts in lung sections. **F.** Kaplan-Meier survival curve comparing *Hif1a*^⁻/⁻^ (n = 9) and wild-type (WT) (n = 11) mice following BLM induction. A 20% loss in body weight was defined as the survival endpoint. **G-J.** Targeted mutation of the *Hif1a* gene mitigates bleomycin-induced pulmonary fibrosis. BLM was intratracheally administered into WT or WT or *Hif1a*^−/−^ mice, and lungs were removed on day 14 after BLM challenge. **G.** Representative images of hematoxylin and eosin (H&E)-stained lung sections for both groups. **H.** Masson Trichrome staining illustrating collagen fibers in mice from both groups. **I.** Quantification of fibrotic area (mm² per view) based on Masson’s trichrome staining, showing a significant reduction in fibrotic area in *Hif1a*^−/−^ mice. **J.** Ashcroft score quantifying fibrosis severity in lung sections, showing a significant reduction in fibrosis in *Hif1a*^−/−^ mice. **K.** Total collagen content in the right lung was quantified using the Sircol assay. WT and *Hif1a*^−/−^ mice were sacrificed on day 14 post-BLM induction, and right lung tissue was collected for analysis. Data are presented as mean ± SEM and are representative of three independent experiments. Each dot represents an individual mouse (n = 13 or 14 per group). Statistical comparison was performed using an unpaired t-test. *p < 0.01.

To evaluate the impact on fibrosis, we compared survival and collagen content in WT and *Hif1a* knockout cohorts 14 days after bleomycin challenge. Survival was improved in *Hif1a* knockout mice relative to WT by Kaplan-Meier analysis (Fig. 4F). Histology showed severe fibrotic changes in WT lungs, including thickened septa, architectural distortion, and inflammatory infiltrates by H&E, whereas *Hif1a* knockout lungs displayed milder abnormalities (Fig. 4G, Fig. S6). Masson’s trichrome staining demonstrated extensive collagen deposition in WT lungs with substantially reduced collagen staining in *Hif1a* knockout lungs (Fig. 4H). Consistently, low-magnification quantification showed a reduced fibrotic area fraction in *Hif1a* knockout mice (Fig. 4I), and Ashcroft scores were significantly lower in *Hif1a* knockout lungs (Fig. 4J). Biochemical collagen quantification using the Sircol soluble collagen assay further corroborated a reduction in right-lung collagen content in *Hif1a* knockout mice compared with WT (Fig. 4K). Collectively, these data indicate that myeloid *Hif1a* deletion mitigates bleomycin-induced pulmonary fibrosis, as reflected by improved survival and concordant reductions across histopathologic, morphometric, and biochemical collagen endpoints.

### Liposomal Echinomycin Inhibits HIF-1α Programs to Reduce Macrophage Burden and Attenuate Bleomycin-Induced Lung Fibrosis

o complement our genetic loss-of-function data and provide an independent pharmacologic test of the HIF-1α hypothesis in macrophage-driven fibrogenesis, we treated bleomycin-challenged WT mice with intranasal liposomal echinomycin (LEM; 0.05 mg/kg once daily for 10 doses starting Day 3 post-bleomycin), a regimen designed to favor lung deposition and minimize systemic exposure. Echinomycin blocks HIF-1-dependent transcription by preventing HIF-1 binding to hypoxia-response elements, and liposomal encapsulation is intended to enhance pulmonary delivery and improve tolerability *in vivo* (33, 34). In bleomycin-challenged mice, LEM treatment was associated with significantly fewer monocyte-derived macrophages/monocytes (MoMp) in BAL and lung tissue compared with vehicle (Fig. 5A, B), while neutrophil counts in BAL and lung were unchanged (Fig. S7), indicating relative selectivity for the macrophage/monocyte lineage. Consistently, BAL TNF-α and IL-6 concentrations were reduced with LEM (Fig. 5C). Immunofluorescence confirmed fewer CD68⁺ cells in LEM-treated lungs (Fig. 5D, E), and co-staining for CD68 and cleaved caspase-3 indicated increased macrophage apoptosis following LEM treatment (Fig. 5F, G), providing a plausible mechanism for the reduced macrophage burden and suggesting diminished pro-fibrotic signaling to stromal cells. In line with this, αSMA immunofluorescence showed robust myofibroblast accumulation in vehicle-treated lungs but substantially lower αSMA signal in LEM-treated mice after bleomycin exposure (Fig. 5H). Histologic analysis at Week 2 demonstrated that LEM markedly attenuated bleomycin-induced fibrosis, with reduced architectural distortion on H&E and decreased collagen deposition on Masson’s trichrome compared with vehicle controls (Fig. 5I-L). Semi-quantitative Ashcroft scoring confirmed a significant reduction in fibrosis severity with LEM (Fig. 5M), and and Sircol-based biochemical measurement of right-lung collagen content showed reduced collagen/ECM accumulation in LEM-treated mice (Fig. 5N). Collectively, these data indicate that pharmacologic HIF-1α inhibition with LEM reduces macrophage/monocyte abundance and inflammatory cytokines, promotes macrophage apoptosis, and attenuates fibrotic remodeling and collagen deposition in bleomycin-injured lungs.

**Fig. 5.**
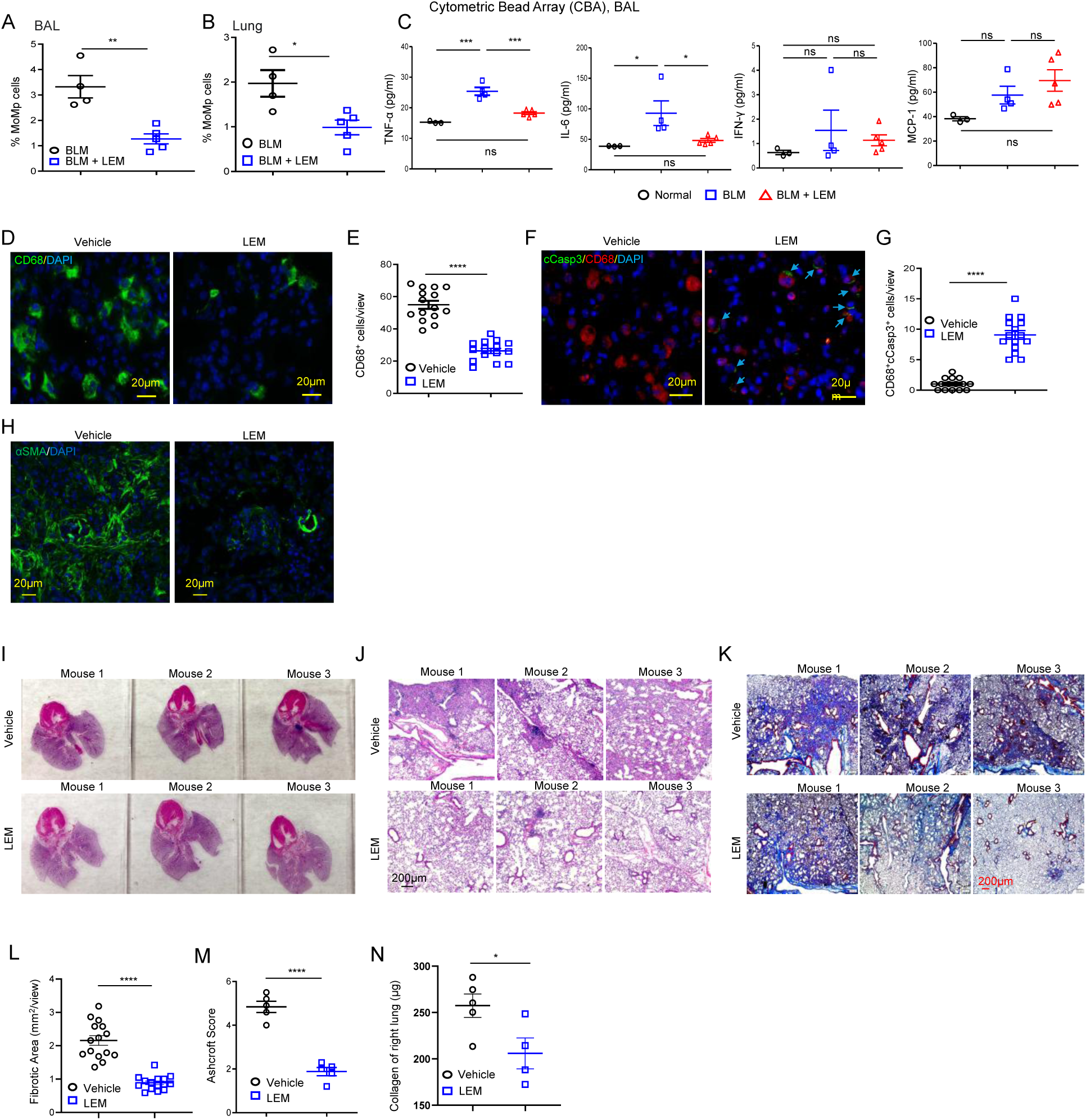
Liposomal Echinomycin Reduces Macrophage Burden and Attenuates Bleomycin-Induced Pulmonary Fibrosis. LEM administration via the intranasal route led to a decrease in the number of macrophages in BLM-induced mice. The mice received intratracheal administration of bleomycin on day 0, followed by intranasal treatment with 0.05 mg/kg of LEM for a cycle of 5 consecutive days with a 2-day interval for the second cycle on day 3. On day 14, bronchoalveolar lavage (BAL) and lung tissues were collected. **A.** Flow cytometry analysis of macrophage frequency in BAL, gated on CD45^+^ subsets from LEM- or vehicle-treated mice. Results are presented as mean ± SEM and are representative of three independent experiments. **B.** Flow cytometry analysis of macrophage frequency in digested lung tissues, gated on CD45^+^ subsets from LEM- or vehicle-treated BLM-induced mice. Summarized data are shown as mean ± SEM for one experiment and are representative of three independent experiments. **C.** Quantification of TNFα, IL-6, IFNγ, and MCP-1 concentrations in the BAL fluid on day 14, determined by cytometric bead array. **D, E.** Immunofluorescence staining of CD68 is shown in (D), with summarized macrophage counts in the tissues presented in (E). Each count is based on three views per lung per mouse at 40X magnification. **F, G.** Co-staining of CD68 and cleaved caspase-3 (cCasp3) is illustrated in (C), while (D) displays the summarized apoptotic macrophage counts for the corresponding tissues at 40X magnification. Each count is based on three views per mouse. Statistical comparisons were made using ANOVA, *P < 0.05, ***P < 0.001, and ****P < 0.0001 versus the indicated group. **H.** Immunofluorescence staining with anti-αSMA antibody to examine αSMA^+^ myofibroblasts in lung sections. **I.** Representative images of whole lung lobes from mice showing reduced fibrotic appearance in LEM-treated mice compared to vehicle-treated controls. **J.** H&E high-power fields illustrate distal parenchymal preservation with LEM compared to dense fibroinflammatory remodeling in vehicle. **K.** Masson’s trichrome staining of lung sections showing collagen deposition. LEM-treated mice display decreased collagen accumulation compared to vehicle-treated controls. **L.** Quantification of fibrotic area (mm²/view) based on Masson’s trichrome staining, showing significantly reduced fibrosis in LEM-treated mice compared to the vehicle group (****p < 0.0001). **M.** Ashcroft score, indicating fibrosis severity, is significantly lower in LEM-treated mice compared to vehicle controls (****p < 0.0001). **N.** Sircol assay-based biochemical quantification of right-lung collagen content (µg) is lower with LEM. Data shown as mean ± SEM; symbols represent individual mice/fields. Statistics: unpaired tests with multiple-comparison corrections where applicable; **** P < 0.0001, * P < 0.05.

### Targeting HIF-1α with sh*Hif1a*-lipid nanoparticles (LNP) attenuate lung fibrosis in BLM-induced PF mice

To further investigate the role of HIF-1α in fibrogenesis and evaluate the therapeutic potential of targeting HIF-1α in BLM-induced PF, we designed lipid nanoparticle (LNP)-encapsulated short hairpin RNA (shRNA) against HIF-1α. The shRNA sequences were previously validated for successful HIF-1α knockdown in a previous study (35, 36). The formulated sh*Hif1a*-LNP exhibited a hydrodynamic diameter of approximately 80.8 nm with high uniformity (PDI ∼0.085), as determined by dynamic light scattering (Fig. 6A). Efficient lung delivery of sh*Hif1a*-LNP was confirmed via intranasal administration, with in vivo tracking using DiD dye and IVIS imaging (Fig. 6B). Mice subjected to BLM-induced fibrosis were treated with sh*Hif1a*-LNP starting from day 3 post-BLM exposure. Histological analysis of lung tissue, including H&E and Masson’s trichrome staining, demonstrated reduced fibrotic remodeling and improved lung architecture in sh*Hif1a*-LNP-treated mice compared to Sc-LNP-treated controls (Fig. 6C, D; Fig. S8). Quantitative assessment of fibrosis severity using the Ashcroft score also indicated a significant attenuation of fibrosis upon sh*Hif1a*-LNP treatment (Fig. 6E). Immunofluorescence further revealed a marked decrease in CD68⁺ macrophage abundance in sh*Hif1a*-LNP-treated lungs (Fig. 6F, G), which was accompanied by reduced αSMA⁺ myofibroblast accumulation in adjacent fibrotic regions (Fig. 6H). Consistent with these histologic and cellular changes, Sircol-based biochemical analysis showed reduced collagen content in the right lung of sh*Hif1a*-LNP-LNP-treated mice compared with Sc-LNP controls (Fig. 6I). Together, these data support the therapeutic potential of sh*Hif1a*-LNP to mitigate BLM-induced lung fibrosis, consistent with a model in which HIF-1α activity sustains a hypoxic, macrophage-rich microenvironment that promotes fibroblast activation and matrix deposition.

**Fig. 6.**
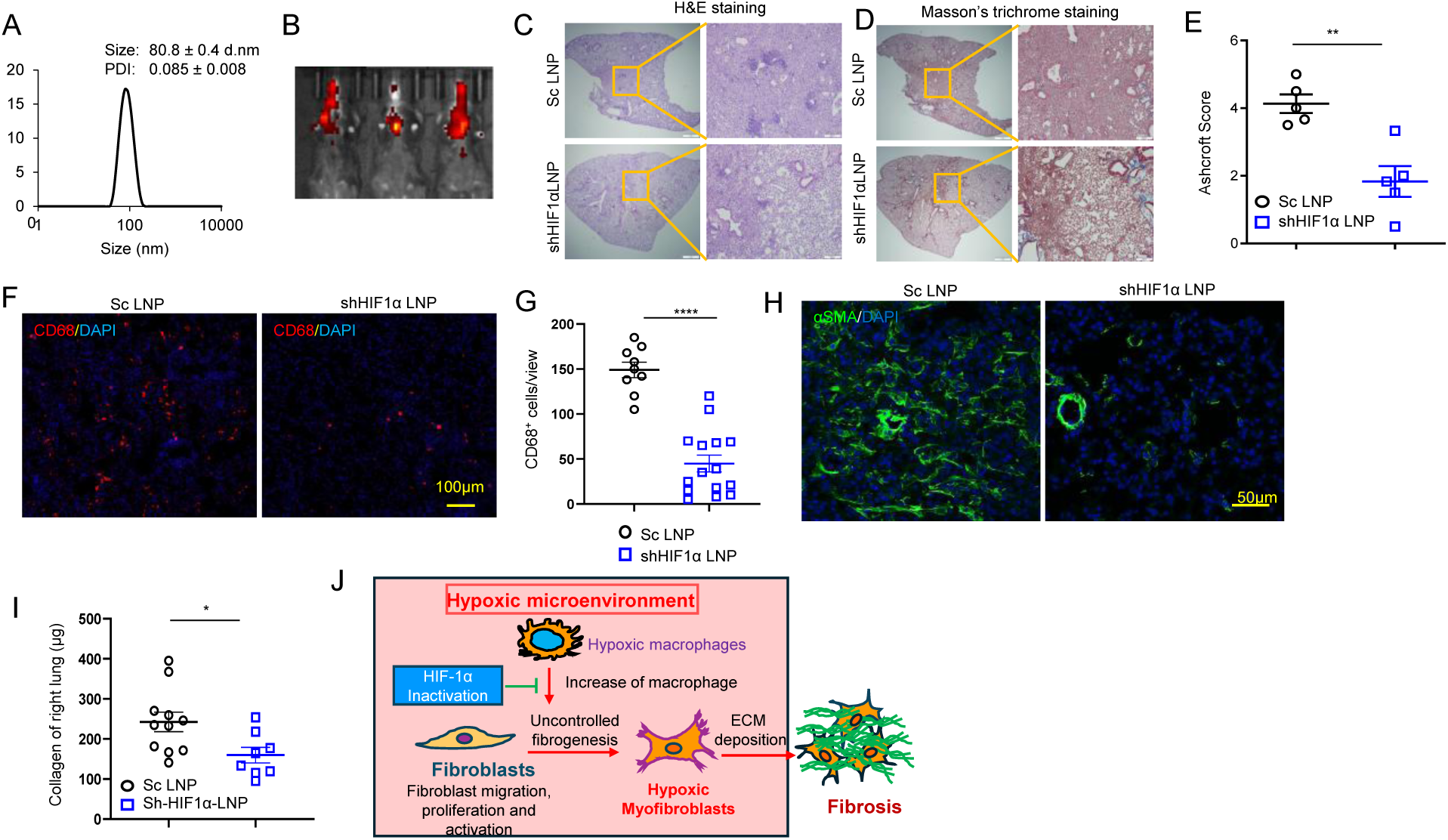
sh*Hif1a*-LNP suppress fibrosis in bleomycin-induced pulmonary fibrosis. **A.** Size distribution of sh*Hif1a*-LNP measured by dynamic light scattering (DLS), including hydrodynamic diameter and polydispersity index (PdI) of sh*Hif1a*-LNP. **B.** Representative in vivo fluorescence images of DiD-labeled sh*Hif1a*-LNP in BLM-induced PF mice 4 h after intranasal administration. sh*Hif1a*-LNP was labeled with the fluorescent dye DiD and delivered by a single intranasal administration to mice with bleomycin-induced pulmonary fibrosis. Fluorescence imaging performed 4 h later demonstrated localization of DiD-sh*Hif1a*-LNP primarily in the lung region. **C,D.** Representative images of H&E (C) and Masson Trichrome (D) stained lung sections in vehicle and sh*Hif1a*-LNP-treated mice. **E.** Ashcroft fibrosis score used for grading fibrosis scale between the vehicle and sh*Hif1a*-LNP groups. **F, G.** Immunofluorescence staining with anti-mouse CD68 shows a reduced number of macrophages in the lungs of BLM-induced mice treated with sh*Hif1a*-LNP compared to control Sr LNP (F). Quantification of macrophages based on CD68 immunofluorescent staining at 20X magnification is summarized (G), with n = 3 views per lung for each mouse. **H.** Immunofluorescence staining with anti-α-SMA antibody to examine α-SMA^+^ myofibroblasts in lung sections. **I.** Total lung collagen content was quantified using the Sircol assay. The right lung of BLM-induced mice treated with either sh*Hif1a*-LNP or control Sr LNP was analyzed for total collagen levels. Treatment with sh*Hif1a*-LNP significantly reduced collagen content in bleomycin-treated lungs compared to control Sr LNP. Data are presented as mean ± SD (sh*Hif1a*-LNP: n = 8; Sr LNP: n = 11) and are representative of two independent experiments. *p < 0.05. **J.** Summary of the underlying mechanism.

Our findings identify a spatially and temporally organized role for macrophages in early fibrogenesis, with HIF-1α acting as a key regulator of the front-zone microenvironment. Across human IPF tissue and mouse time-course studies, HIF-1α activation marks a subset of macrophages positioned at advancing fibrotic fronts, where they align with hypoxia, chemokine cues, and early fibroblast activation, rather than within mature scar. In this setting, HIF-1α-positive macrophages associate with fibroblast activation, followed by fibroblast migration, proliferation, and differentiation into αSMA⁺ myofibroblasts that drive extracellular matrix deposition and lesion expansion. The coordinated presence of hypoxic macrophages, activated fibroblasts, and αSMA⁺ myofibroblasts at the front highlights a previously underappreciated compartment in which HIF-1α activity shapes the microenvironment of progression (Fig. 6J). The novelty of our study lies in defining this spatial-temporal ordering at fibrotic fronts, validating it in both human and mouse lung environments, and showing with complementary genetic, pharmacologic, and RNA-interference approaches that interrupting myeloid HIF-1α reduces macrophage accumulation, fibroblast activation, and downstream matrix deposition.

## Discussion

Pulmonary fibrosis reflects dynamic interactions between immune and mesenchymal compartments within evolving tissue niches (19, 37). In progressive pulmonary fibrosis, current antifibrotics slow but rarely halt decline, highlighting the need to define mechanism-based, compartment-specific drivers of lesion propagation (2-4, 30). Although hypoxia/HIF-1α signaling has long been linked to fibrotic remodeling, prior work has often been correlative or focused on fibroblast-intrinsic pathways, leaving unresolved where and when HIF-1α acts to sustain lesion expansion (19, 38-42). Here, integrating human IPF tissue and datasets, single-cell transcriptomics, and complementary in vivo perturbations, we identify a convergent front-zone program in which HIF-1α activity is enriched in macrophage- and fibroblast-dense niches, hypoxic macrophages concentrate at advancing fronts, and disruption of *Hif1a* reduces macrophage accumulation, fibroblast activation, and matrix deposition (19, 38-45). These findings support a front-zone hypoxia-myeloid HIF-1α axis consistent with emerging spatiotemporal frameworks emphasizing niche-localized macrophage programs in fibrotic progression (7).

Multiple studies report increased HIF-1α-associated signatures in IPF tissue and biofluids, with higher scores linked to worse outcomes (19, 38-40, 42, 43), and *HIF1A* has been implicated in reinforcing profibrotic fibroblast programs (40). Consistent with this, our LGRC analysis confirms increased *HIF1A* expression in IPF and association with GAP score (38, 40, 43). Importantly, our data place HIF-1α activation into a compartment-resolved framework: multiplex imaging identifies CD68⁺ macrophages as a prominent HIF-1α⁺ source in fibrotic regions, while scRNA-seq shows enrichment of HIF-1α-linked programs in monocyte/macrophage and fibroblast clusters, including growth factors, chemokines, and matrix-regulatory pathways implicated in macrophage-fibroblast crosstalk (6, 37, 38, 44-47).

Our findings align with prior single-cell studies identifying profibrotic macrophage populations in IPF, including SPP1^hi^/MERTK⁺ macrophages, ApoE⁺ monocyte-derived alveolar macrophages, and TREM2-associated macrophage survival/profibrotic activity (28, 48, 49). Extending these concepts spatially, we show that macrophages are enriched at lesion margins while mature collagen-dense cores are relatively macrophage-poor in both mouse and human lungs (6, 19, 37, 44, 45, 50). In bleomycin injury, macrophage expansion occurs early-to-mid, whereas PDGFRα⁺ fibroblasts and αSMA⁺ myofibroblasts rise later, with HIF-1α detectable in both compartments (Fig. 2) (47, 51-53). Serial sections show hypoxic rims bordering nascent αSMA⁺ foci that are enriched for CD68⁺ macrophages, consistent with a hypoxic myeloid “cuff” at the advancing front (6, 37, 44, 45). This front-enrichment helps reconcile why consolidated scar can appear macrophage-low despite strong evidence that macrophages are required for fibrogenesis: key myeloid effector programs may be concentrated at the leading edge where remodeling cues are maximal (7, 37, 46, 47).

Because bleomycin inflammation is often described as peaking around Days 4–7 with partial contraction thereafter (54, 55), we note that our leukocyte kinetics were quantified as absolute counts from whole-lung digests, which may differ from histopathologic impressions from thin sections. In our dataset, T and B lymphocytes were highest at Days 10-14 (Fig. S2), consistent with time-resolved cytometry reporting late increases in adaptive compartments, including a Day 14 peak in CD19⁺ B cells (26), and with the concept that spatially restricted immune aggregates in IPF/UIP can be under-sampled by routine histology (56-59). These observations support interpreting late lymphocyte expansion as a remodeling-associated feature rather than persistence of the initial acute inflammatory surge.

HIF-1α has context-dependent roles in lung injury/repair, with protective effects in some epithelial/endothelial settings and profibrotic effects in mesenchymal compartments (39, 41-43). In myeloid cells, hypoxia-regulated pathways (including upstream regulators such as VHL) modulate inflammation and fibrosis, but direct testing of myeloid *Hif1a* loss during fibrotic progression has been limited (51, 60). Using LysM-Cre conditional knockout, we show that myeloid HIF-1α is required for full expression of bleomycin-induced fibrosis: *Hif1a* deletion reduces monocyte-derived and alveolar macrophage populations, decreases CD68⁺ density at the parenchyma-fibrotic interface, and attenuates collagen deposition and Ashcroft scores, with minimal effect on neutrophil numbers (47, 51-53, 61, 62). Thus, beyond bulk signatures or fibroblast-intrinsic circuits (16, 30-34, 37), our data establish myeloid HIF-1α as a causal determinant of macrophage persistence at fibrotic fronts and downstream matrix accumulation in vivo.

Two lung-directed interventions converged with genetics. Inhaled liposomal echinomycin reduced lung/BAL monocyte-macrophage numbers, increased macrophage apoptosis, decreased inflammatory mediators, and attenuated fibrosis and collagen content; given echinomycin’s systemic limitations, these findings support a lung-local strategy distinct from systemic HIF blockade (33, 63-65). In parallel, intranasal sh*Hif1a*-LNP achieved robust lung deposition and reduced CD68⁺ macrophages, αSMA⁺ myofibroblasts, and collagen deposition (66, 67). Together, myeloid deletion, inhaled echinomycin, and inhaled sh*Hif1a*-LNP provide multimodal evidence that front-zone myeloid HIF-1α is both necessary and therapeutically addressable in pulmonary fibrosis.

Mechanistically, our data support a model in which hypoxic, HIF-1α⁺ macrophages at fibrotic fronts promote fibroblast activation and matrix remodeling via growth factors, cytokines, and chemokines, while fibroblasts activate HIF-1α-linked matrix programs (38, 40-43, 47). These findings nominate a lung-local, macrophage-enriched HIF-1α strategy as a potential adjunct to antifibrotics, enabling future testing of combination regimens using standard endpoints (2-4, 51). Timing may be critical, as inhaled interventions were initiated after inflammatory onset but before maximal consolidation, when macrophage fronts and nascent αSMA⁺ islands are prominent (47, 51-53), potentially limiting systemic exposure (33, 63, 64, 66, 67).

Limitations include reliance on bleomycin as an acute injury model (47, 51-53), broad LysM-Cre targeting and variable recombination across subsets (61), and potential effects of inhaled interventions on non-myeloid HIF-1α-expressing cells (61, 64, 66, 67). Translation will also need to account for patient heterogeneity and integration with standard antifibrotic therapy (20, 51). Despite these caveats, our study supports a spatially and temporally organized front-zone program in which myeloid HIF-1α sustains macrophage persistence and drives fibrotic remodeling, motivating follow-up studies to refine therapeutic windows, define the key macrophage subsets, and test lung-directed HIF-1α modulation in IPF and related ILDs (42, 51, 63, 64, 67).

## Author contributions

YW, YL, and WD designed the study, analyzed the data, supervised the research, and wrote the manuscript.YL and YW designed and conducted the research and contributed to manuscript preparation. CB and YW performed animal experiments and flow cytometry, assisting in result interpretation. YYW and AS conducted histological experiments and contributed to data analysis. YL carried out molecular and biological experiments, while PZ conducted bioinformatics analyses. JP, DL and FDM were responsible for sample collection and processing.

## Acknowledgments

This research was supported by the National Institutes of Health (NIH) through grants CA219150 and CA227671 (to Y.W.) and HL157533, The Dreiling Foundation and HL24127301 (to W.D.). Additional support was provided by Maryland’s Cancer Moonshot Initiative in Pediatric Cancer Research and the University of Maryland, Baltimore Institute for Clinical & Translational Research (to Y.W.). We would also like to thank the sarcoidosis and IPF patients for study participation.

## Materials and methods

### Mice

All procedures involving experimental animals were approved by Institutional Animal Care and Use Committees (IACUC) of the University of Maryland School of Medicine where this work was performed. All animal experiments were conducted according to guidelines established by the NIH Guide for the Care and Use of Laboratory Animals (National Academies Press, 2011). *Hif1a^flox/flox^*and LysM^Cre^ mice were purchased from the Jackson Laboratory. C57BL/6 mice were purchased from Charles River Laboratories. Male and female mice were used for all studies. Animals were group housed and maintained under standard conditions (temperature, humidity, and light controlled, and standard diet) for mouse research at the Research Animal Facilities at University of Maryland School of Medicine. None of the animals selected for experiments had received prior treatment or were involved in any previous procedures.

### Single-cell RNA-seq analysis

Single-cell RNA-seq count matrices from Adams et al. (Single-cell RNA-seq reveals ectopic and aberrant lung-resident cell populations in idiopathic pulmonary fibrosis) were downloaded from the Gene Expression Omnibus (GEO) database (accession number GSE136831) (16), and processed in R (v4.1.3) using Seurat (v5). The count matrices were loaded using Read10X, and batch effects were corrected via Reciprocal Principal Component Analysis (RPCA) before normalization with the LogNormalization method. Genes detected in fewer than 3 cells were excluded, and cells with fewer than 200 genes or more than 10,000 genes were removed. Additionally, cells with >10% unique molecular identifiers (UMIs) mapped to mitochondrial or erythrocyte genes were discarded. UMI counts were scaled by library size, multiplied by 10,000, and log-transformed. Highly variable genes were identified using FindVariableFeatures, and principal component analysis (PCA) was performed, selecting the top components based on PCElbowPlot. Dimensionality reduction and clustering were conducted at resolution = 0.9, followed by visualization using t-distributed stochastic neighbor embedding (t-SNE). Marker genes were identified using the Bimod test in FindAllMarkers with adjusted P-value < 0.01, log2 fold change ≥ 0.25, and genes detected in fewer than 10% of cluster cells excluded. All samples were normalized via sctransform and integrated using RPCA in IntegrateData. The top 2,000 highly variable genes and first 30 principal components (PCs) were used for alignment before clustering and Uniform Manifold Approximation and Projection (UMAP) visualization. Genes with log2 average expression difference ≥ 0.585 and P < 0.05 were classified as marker genes. Cell clusters were annotated using canonical markers of known cell types, and the expression of HIF1A and its regulated genes across clusters was visualized using FeaturePlot and DotPlot in Seurat.

### Bleomycin-Induced Lung Injury and Fibrosis, and therapeutic agents

For each experiment, age- and weight-matched groups of mice were used. Mice were anesthetized with intraperitoneal pentobarbital, their lungs were intubated orally with a 20-gauge Angiocath (Franklin Lanes, NJ), and two 50-μl sterile aliquots of PBS (control) or 0.025 IU bleomycin (AAP Pharmaceuticals LLC, Shamburg, IL) were instilled through the catheter, 3 minutes apart. After each aliquot, the mice were placed on their right side and then left side for 10–15 seconds. Mice were euthanized at 14 or 21 days after the instillation of bleomycin. Bleomycin was purchased from Sigma. Echinomycin was formulated with liposomes as previously described (35).

### Flow cytometry

Inflammatory leukocytes in bronchoalveolar lavage fluid and collagenase digests of lung tissue. Mice were euthanized, and the trachea was cannulated using an 18-gauge needle. The lungs underwent two lavages with PBS, and the collected lavage fluid was pooled, centrifuged, and the supernatants were then collected and stored at -80°C until CBA assay. The resulting cell pellet was resuspended and washed with PBS and subjected to staining for flow cytometry analysis. Simultaneously, following lavage, the lung vascular bed was perfused with PBS, and the lungs were excised, minced, and digested using a digesting buffer containing 1 mg/ml collagenase, and 30 U/ml DNase at 37°C for 30 minutes. The resulting suspension was dispersed through repeated aspiration with a pipette, and the red blood cells were lysed using BD FACS Lysing Solution. The cells were filtered through a 100-µm nylon mesh, stained with antibodies, and subjected to flow cytometry analysis. Antibodies used were PerCP-Cy5.5-mGr-1, PE-mCD64, PE-Cy5-mCD24, APC-mMHC-II (BioLegend). BUV and BV series antibodies used for Cytek Aurora included the following (all from BD Bioscience, San Jose, CA): BUV496-mCD45, AF700-mSiglecF, BV605-mCD11c,, and BV480-mCD11b. The stained cells were analyzed using the BD FACS Canto II or Cytek Aurora flow cytometers.

Flow Cytometry For surface staining, cells were first washed in 1xPBS, resuspended in 1xPBS with 1x Live/Dead™ Fixable Aqua Dead Cell Stain (Life Technologies, Frederick, MD), and left on ice for 5 minutes to label dead cells. Then, the cells were washed in SB, and stained for 30 minutes on ice with antibodies at 1:100 dilution in SB. In certain cases, intracellular staining of IFNγ (1:100 dilution) was further carried out on surface-stained cells with eBioscience™ Foxp3/Transcription Factor Staining Buffer Set (Life Technologies, Frederick, MD), according to the manufacturer’s instructions. All data was collected on a BD FACS Canto II flow cytometer and subsequently analyzed using FlowJo v10.6.1 software.

### Histology and Immunofluorescence

Tissues harvested from the mice were fixed in 10% neutral-buffered formalin before embedding in paraffin blocks and cutting into 4-μm sections. Hematoxylin and eosin (H&E) and MTC (Masson’s Trichrome Kit, Sigma-Aldrich, St Louis, MO, USA) were stained according to manufacturers protocol. Fibrotic scoring was performed by a blinded pathologist according to the scale with a grade from 0 to 8 using the modified Ashcroft Scale described in the references (68, 69). Pathologist scoring was performed using a 20-fold objective optimized for histological assessment of lung fibrosis.

Additional unstained slides were returned for immunofluorescence analysis. After deparaffinization and rehydration with xylene and ethanol, tissue sections were treated with 10 mM sodium citrate buffer, pH 6.0. The sections were permeabilized with 0.3% Triton X-100 in 10 mM Tris-HCl buffer for 30 min. After blocking with 2% bovine serum albumin (BSA) for 60 min, sections were incubated with primary antibody diluted in 10 mM Tris-HCl buffer containing 2% BSA at 4°C, overnight, with subsequent staining with secondary antibody in BSA-Tris-HCl buffer at room temperature for 2-4 h. The nuclei were stained with DAPI. Slides were mounted with Prolong Antifade mounting buffer (Invitrogen, Carlsbad, CA 92008). For hypoxic probe staining, mice received hypoxyprobe (pimonidazole) 90 minutes prior to euthanasia according to the manufacturer’s instructions. The lung tissues were then frozen and sectioned and stained with the hypoxyprobe-1 for anti-pimonidazole (Probe, green) and anti-mouse CD68, or αSMA, counterstained with DAPI. Antibodies for mCD68 (FA-11, abcam), mCD206 (2A6A10, proteintech), cleaved caspase-3 (Asp175, 5A1E), αSMA (abcam), Collage I (Abcam), TGFβ (Abcam), cMYC (Cell signaling Technology) and VEGF (Santa Cruz) were used for immunofluorescence. HIF-1α antibody (GeneTex) was used for immunochemistry Staining. After mounting, the slides were analyzed with an Olympus BX53 Light Microscope equipped with DP80 Dual CCD camera and CellSens Imaging software.

### Sircol Assay for Collagen Quantification

The collagen content in lung tissue was measured using the Sircol Collagen Assay (BioColor Ltd., Newton Abbey, UK) following the manufacturer’s protocol. Briefly, the right lobes of the lung were homogenized in acid-pepsin solution and incubated overnight at 4°C to facilitate collagen extraction. After neutralization and isolation, the purified collagen was obtained. To quantify collagen, Sircol dye reagent was added to the extracted collagen, forming a collagen-dye complex, which subsequently precipitated. The precipitate was then dissolved using an alkali reagent, and the absorbance was measured at 556 nm using a spectrophotometer. The collagen concentration was determined by comparison with a standard curve and expressed as µg of collagen per right lung homogenate.

### LNP formulation and siRNA encapsulation

The ionizable lipid MC3 (Broadpharm), DSPC, and cholesterol (Avanti, AL) were used for LNP formulation. Lipophilic dyes, 3,3′-dioctadecyloxacarbo cyanine perchlorate (DiOC18), were obtained from Invitrogen (Carlsbad, CA). Lipids were mixed at molar ratios (mol/mol) of DSPC/Chol/MC3/PEG-DMG/DiOC18 (10/37.5/50/1.5/1). LNPs were synthesized at a 3:1 aqueous-to-organic phase flow rate ratio (FRR). The aqueous phase contained siRNA (300 µg/mL) in 50 mM acetate buffer (pH 4.0), and the lipid phase was prepared in ethanol. The final lipid concentration was 31 mM. Following synthesis, LNPs were diluted in DPBS (1X, pH 7.0) and purified using 100 kDa MWCO Amicon centrifugal filters.

LNP characterization was performed via dynamic light scattering (DLS) (Zetasizer Nano ZS90, Malvern Panalytical, UK). Samples were diluted 1:1000 in DPBS (1X). Three measurements were obtained per sample, and mean values with standard deviations were calculated.

### Measurement of BAL Cytokines

Cytokine levels in BAL fluid were measured using the Becton Dickinson (BD) Cytometric Bead Array (CBA) Mouse Inflammation Kit (BD Biosciences, San Jose, CA) following the manufacturer’s protocol. The concentrations of IFN-γ, IL-6, TNF-α, and MCP-1 were quantified using flow cytometry. Standard curves for each cytokine were generated using the reference cytokine concentrations provided in the kit.

### Statistics

All experiments were replicated at least twice with similar results. Appropriate statistical tests were selected on the basis of whether the data with outlier deletion was normally distributed by using the D’Agostino & Pearson normality test. Data comparing 2 groups were analyzed by unpaired, 2-tailed Student’s t test. Unless otherwise noted in the figure legends, 1-way analysis of variance (ANOVA) with Sidak’s post hoc test was used for multiple comparisons, and 2-way ANOVA for analysis of tumor kinetics. The correlation coefficient and P value for linear regression were calculated by Pearson’s method. Sample sizes were chosen with adequate statistical power on the basis of the literature and past experience. In the graphs, data are shown as mean ± SEM, indicated by horizontal line and y-axis error bars, respectively. Statistical calculations were performed using Prism 8 software (GraphPad Software). NS in the figures indicates no significant difference. A P value of less than 0.05 was considered significant: *P < 0.05, **P < 0.01, ***P < 0.001, ****P < 0.0001.

## References

1. Richeldi L, Collard HR, Jones MG. Idiopathic pulmonary fibrosis. Lancet 2017;389:1941–1952.

2. Faverio P, Piluso M, De Giacomi F, Della Zoppa M, Cassandro R, Harari S, Luppi F, Pesci A. Progressive fibrosing interstitial lung diseases: Prevalence and characterization in two italian referral centers. Respiration; international review of thoracic diseases 2020;99:838–845.

3. Chianese M, Screm G, Salton F, Confalonieri P, Trotta L, Barbieri M, Ruggero L, Mari M, Reccardini N, Geri P, Hughes M, Lerda S, Confalonieri M, Mondini L, Ruaro B. Pirfenidone and nintedanib in pulmonary fibrosis: Lights and shadows. Pharmaceuticals 2024;17.

4. Trachalaki A, Sultana N, Wells AU. An update on current and emerging drug treatments for idiopathic pulmonary fibrosis. Expert opinion on pharmacotherapy 2023;24:1125–1142.

5. Herrera JA, Dingle L, Montero MA, Venkateswaran RV, Blaikley JF, Lawless C, Schwartz MA. The uip/ipf fibroblastic focus is a collagen biosynthesis factory embedded in a distinct extracellular matrix. JCI insight 2022;7.

6. Franzen L, Olsson Lindvall M, Huhn M, Ptasinski V, Setyo L, Keith BP, Collin A, Oag S, Volckaert T, Borde A, Lundeberg J, Lindgren J, Belfield G, Jackson S, Ollerstam A, Stamou M, Stahl PL, Hornberg JJ. Mapping spatially resolved transcriptomes in human and mouse pulmonary fibrosis. Nature genetics 2024;56:1725–1736.

7. Behmoaras J, Mulder K, Ginhoux F, Petretto E. The spatial and temporal activation of macrophages during fibrosis. Nature reviews Immunology 2025;25:816–830.

8. Lin N, Simon MC. Hypoxia-inducible factors: Key regulators of myeloid cells during inflammation. The Journal of clinical investigation 2016;126:3661–3671.

9. Brereton CJ, Yao L, Davies ER, Zhou Y, Vukmirovic M, Bell JA, Wang S, Ridley RA, Dean LSN, Andriotis OG, Conforti F, Brewitz L, Mohammed S, Wallis T, Tavassoli A, Ewing RM, Alzetani A, Marshall BG, Fletcher SV, Thurner PJ, Fabre A, Kaminski N, Richeldi L, Bhaskar A, Schofield CJ, Loxham M, Davies DE, Wang Y, Jones MG. Pseudohypoxic hif pathway activation dysregulates collagen structure-function in human lung fibrosis. eLife 2022;11.

10. Stothers CL, Luan L, Fensterheim BA, Bohannon JK. Hypoxia-inducible factor-1alpha regulation of myeloid cells. Journal of molecular medicine 2018;96:1293–1306.

11. Stancil IT, Michalski JE, Hennessy CE, Hatakka KL, Yang IV, Kurche JS, Rincon M, Schwartz DA. Interleukin-6-dependent epithelial fluidization initiates fibrotic lung remodeling. Sci Transl Med 2022;14:eabo5254.

12. Mia MM, Ghani S, Cibi DM, Bogireddi H, Nilanthi U, Selvan A, Wong WSF, Singh MK. Yap/taz are crucial regulators of macrophage-mediated pulmonary inflammation and fibrosis after bleomycin-induced injury. The European respiratory journal 2025;65.

13. Tanner L, Single AB, Bhongir RKV, Heusel M, Mohanty T, Karlsson CAQ, Pan L, Clausson CM, Bergwik J, Wang K, Andersson CK, Oommen RM, Erjefalt JS, Malmstrom J, Wallner O, Boldogh I, Helleday T, Kalderen C, Egesten A. Small-molecule-mediated ogg1 inhibition attenuates pulmonary inflammation and lung fibrosis in a murine lung fibrosis model. Nature communications 2023;14:643.

14. Chen K, Han H, Zhao S, Xu B, Yin B, Lawanprasert A, Trinidad M, Burgstone BW, Murthy N, Doudna JA. Lung and liver editing by lipid nanoparticle delivery of a stable crispr-cas9 ribonucleoprotein. Nature biotechnology 2025;43:1445–1457.

15. Zhu M, Huang Y, Bender ME, Girard L, Kollipara R, Eglenen-Polat B, Naito Y, Savage TK, Huffman KE, Koyama S, Kumanogoh A, Minna JD, Johnson JE, Akbay EA. Evasion of innate immunity contributes to small cell lung cancer progression and metastasis. Cancer research 2021;81:1813–1826.

16. Aviles P, Altares R, van Andel L, Lubomirov R, Fudio S, Rosing H, Marquez Del Pino FM, Tibben MM, Benedit G, Nan-Offeringa L, Luepke Estefan XE, Francesch A, Zeaiter A, Cuevas C, Schellens JHM, Beijnen JH. Metabolic disposition of lurbinectedin, a potent selective inhibitor of active transcription of protein-coding genes, in nonclinical species and patients. Drug Metab Dispos 2022;50:327–340.

17. Copple BL, Kaska S, Wentling C. Hypoxia-inducible factor activation in myeloid cells contributes to the development of liver fibrosis in cholestatic mice. The Journal of pharmacology and experimental therapeutics 2012;341:307–316.

18. Karakiulakis G, Papakonstantinou E, Aletras AJ, Tamm M, Roth M. Cell type-specific effect of hypoxia and platelet-derived growth factor-bb on extracellular matrix turnover and its consequences for lung remodeling. The Journal of biological chemistry 2007;282:908–915.

19. Tzouvelekis A, Harokopos V, Paparountas T, Oikonomou N, Chatziioannou A, Vilaras G, Tsiambas E, Karameris A, Bouros D, Aidinis V. Comparative expression profiling in pulmonary fibrosis suggests a role of hypoxia-inducible factor-1alpha in disease pathogenesis. American journal of respiratory and critical care medicine 2007;176:1108–1119.

20. Ruthenborg RJ, Ban JJ, Wazir A, Takeda N, Kim JW. Regulation of wound healing and fibrosis by hypoxia and hypoxia-inducible factor-1. Molecules and cells 2014;37:637–643.

21. Lee JW, Ko J, Ju C, Eltzschig HK. Hypoxia signaling in human diseases and therapeutic targets. Experimental & molecular medicine 2019;51:1–13.

22. Mosser DM, Edwards JP. Exploring the full spectrum of macrophage activation. Nature reviews Immunology 2008;8:958–969.

23. Wynn TA, Ramalingam TR. Mechanisms of fibrosis: Therapeutic translation for fibrotic disease. Nature medicine 2012;18:1028–1040.

24. Clara CA, Marie SK, de Almeida JR, Wakamatsu A, Oba-Shinjo SM, Uno M, Neville M, Rosemberg S. Angiogenesis and expression of pdgf-c, vegf, cd105 and hif-1alpha in human glioblastoma. Neuropathology : official journal of the Japanese Society of Neuropathology 2014;34:343-352.

25. Saddouk FZ, Kuzemczak A, Saito J, Greif DM. Endothelial hifalpha/pdgf-b to smooth muscle beclin1 signaling sustains pathological muscularization in pulmonary hypertension. JCI insight 2024;9.

26. Bordag N, Biasin V, Schnoegl D, Valzano F, Jandl K, Nagy BM, Sharma N, Wygrecka M, Kwapiszewska G, Marsh LM. Machine learning analysis of the bleomycin mouse model reveals the compartmental and temporal inflammatory pulmonary fingerprint. iScience 2020;23:101819.

27. Misharin AV, Morales-Nebreda L, Reyfman PA, Cuda CM, Walter JM, McQuattie-Pimentel AC, Chen CI, Anekalla KR, Joshi N, Williams KJN, Abdala-Valencia H, Yacoub TJ, Chi M, Chiu S, Gonzalez-Gonzalez FJ, Gates K, Lam AP, Nicholson TT, Homan PJ, Soberanes S, Dominguez S, Morgan VK, Saber R, Shaffer A, Hinchcliff M, Marshall SA, Bharat A, Berdnikovs S, Bhorade SM, Bartom ET, Morimoto RI, Balch WE, Sznajder JI, Chandel NS, Mutlu GM, Jain M, Gottardi CJ, Singer BD, Ridge KM, Bagheri N, Shilatifard A, Budinger GRS, Perlman H. Monocyte-derived alveolar macrophages drive lung fibrosis and persist in the lung over the life span. J Exp Med 2017;214:2387–2404.

28. Morse C, Tabib T, Sembrat J, Buschur KL, Bittar HT, Valenzi E, Jiang Y, Kass DJ, Gibson K, Chen W, Mora A, Benos PV, Rojas M, Lafyatis R. Proliferating spp1/mertk-expressing macrophages in idiopathic pulmonary fibrosis. The European respiratory journal 2019;54.

29. Bailey JI, Puritz CH, Senkow KJ, Markov NS, Diaz E, Jonasson E, Yu Z, Swaminathan S, Lu Z, Fenske S, Grant RA, Abdala-Valencia H, Mylvaganam RJ, Ludwig A, Miller J, Cumming RI, Tighe RM, Gowdy KM, Kalhan R, Jain M, Bharat A, Kurihara C, San Jose Estepar R, San Jose Estepar R, Washko GR, Shilatifard A, Sznajder JI, Ridge KM, Budinger GRS, Braun R, Misharin AV, Sala MA. Profibrotic monocyte-derived alveolar macrophages are expanded in patients with persistent respiratory symptoms and radiographic abnormalities after covid-19. Nat Immunol 2024;25:2097–2109.

30. Raghu G, Remy-Jardin M, Richeldi L, Thomson CC, Inoue Y, Johkoh T, Kreuter M, Lynch DA, Maher TM, Martinez FJ, Molina-Molina M, Myers JL, Nicholson AG, Ryerson CJ, Strek ME, Troy LK, Wijsenbeek M, Mammen MJ, Hossain T, Bissell BD, Herman DD, Hon SM, Kheir F, Khor YH, Macrea M, Antoniou KM, Bouros D, Buendia-Roldan I, Caro F, Crestani B, Ho L, Morisset J, Olson AL, Podolanczuk A, Poletti V, Selman M, Ewing T, Jones S, Knight SL, Ghazipura M, Wilson KC. Idiopathic pulmonary fibrosis (an update) and progressive pulmonary fibrosis in adults: An official ats/ers/jrs/alat clinical practice guideline. American journal of respiratory and critical care medicine 2022;205:e18–e47.

31. Nabet BY, Hamidi H, Lee MC, Banchereau R, Morris S, Adler L, Gayevskiy V, Elhossiny AM, Srivastava MK, Patil NS, Smith KA, Jesudason R, Chan C, Chang PS, Fernandez M, Rost S, McGinnis LM, Koeppen H, Gay CM, Minna JD, Heymach JV, Chan JM, Rudin CM, Byers LA, Liu SV, Reck M, Shames DS. Immune heterogeneity in small-cell lung cancer and vulnerability to immune checkpoint blockade. Cancer Cell 2024;42:429–443 e424.

32. Tang J, Wang T, Wu H, Bao X, Xu K, Ren T. Efficacy and toxicity of lurbinectedin in subsequent systemic therapy of extensive-stage small cell lung cancer: A meta-analysis. BMC cancer 2024;24:1351.

33. Bailey CM, Liu Y, Peng G, Zhang H, He M, Sun D, Zheng P, Liu Y, Wang Y. Liposomal formulation of hif-1alpha inhibitor echinomycin eliminates established metastases of triple-negative breast cancer. Nanomedicine : nanotechnology, biology, and medicine 2020;29:102278.

34. Wang Y, Liu Y, Bailey C, Zhang H, He M, Sun D, Zhang P, Parkin B, Baer MR, Zheng P, Malek SN, Liu Y. Therapeutic targeting of tp53-mutated acute myeloid leukemia by inhibiting hif-1alpha with echinomycin. Oncogene 2020;39:3015–3027.

35. Liu Y, Nelson MV, Bailey C, Zhang P, Zheng P, Dome JS, Liu Y, Wang Y. Targeting the hif-1alpha-igfbp2 axis therapeutically reduces igf1-akt signaling and blocks the growth and metastasis of relapsed anaplastic wilms tumor. Oncogene 2021;40:4809–4819.

36. Peng G, Wang Y, Ge P, Bailey C, Zhang P, Zhang D, Meng Z, Qi C, Chen Q, Chen J, Niu J, Zheng P, Liu Y, Liu Y. The hif1alpha-pdgfd-pdgfralpha axis controls glioblastoma growth at normoxia/mild-hypoxia and confers sensitivity to targeted therapy by echinomycin. Journal of experimental & clinical cancer research : CR 2021;40:278.

37. Zhang L, Wang Y, Wu G, Xiong W, Gu W, Wang CY. Macrophages: Friend or foe in idiopathic pulmonary fibrosis? Respiratory research 2018;19:170.

38. Aquino-Galvez A, Gonzalez-Avila G, Jimenez-Sanchez LL, Maldonado-Martinez HA, Cisneros J, Toscano-Marquez F, Castillejos-Lopez M, Torres-Espindola LM, Velazquez-Cruz R, Rodriguez VHO, Flores-Soto E, Solis-Chagoyan H, Cabello C, Zuniga J, Romero Y. Dysregulated expression of hypoxia-inducible factors augments myofibroblasts differentiation in idiopathic pulmonary fibrosis. Respiratory research 2019;20:130.

39. Delbrel E, Soumare A, Naguez A, Label R, Bernard O, Bruhat A, Fafournoux P, Tremblais G, Marchant D, Gille T, Bernaudin JF, Callard P, Kambouchner M, Martinod E, Valeyre D, Uzunhan Y, Planes C, Boncoeur E. Hif-1alpha triggers er stress and chop-mediated apoptosis in alveolar epithelial cells, a key event in pulmonary fibrosis. Scientific reports 2018;8:17939.

40. Epstein Shochet G, Bardenstein-Wald B, McElroy M, Kukuy A, Surber M, Edelstein E, Pertzov B, Kramer MR, Shitrit D. Hypoxia inducible factor 1a supports a pro-fibrotic phenotype loop in idiopathic pulmonary fibrosis. International journal of molecular sciences 2021;22.

41. Romero Y, Balderas-Martinez YI, Vargas-Morales MA, Castillejos-Lopez M, Vazquez-Perez JA, Calyeca J, Torres-Espindola LM, Patino N, Camarena A, Carlos-Reyes A, Flores-Soto E, Leon-Reyes G, Sierra-Vargas MP, Herrera I, Luis-Garcia ER, Ruiz V, Velazquez-Cruz R, Aquino-Galvez A. Effect of hypoxia in the transcriptomic profile of lung fibroblasts from idiopathic pulmonary fibrosis. Cells 2022;11.

42. Yang L, Gilbertsen A, Xia H, Benyumov A, Smith K, Herrera J, Racila E, Bitterman PB, Henke CA. Hypoxia enhances ipf mesenchymal progenitor cell fibrogenicity via the lactate/gpr81/hif1alpha pathway. JCI insight 2023;8.

43. Nho RS, Rice C, Prasad J, Bone H, Farkas L, Rojas M, Horowitz JC. Persistent hypoxia promotes myofibroblast differentiation via gpr-81 and differential regulation of ldh isoenzymes in normal and idiopathic pulmonary fibrosis fibroblasts. Physiological reports 2023;11:e15759.

44. Adams TS, Schupp JC, Poli S, Ayaub EA, Neumark N, Ahangari F, Chu SG, Raby BA, DeIuliis G, Januszyk M, Duan Q, Arnett HA, Siddiqui A, Washko GR, Homer R, Yan X, Rosas IO, Kaminski N. Single-cell rna-seq reveals ectopic and aberrant lung-resident cell populations in idiopathic pulmonary fibrosis. Science advances 2020;6:eaba1983.

45. Reyfman PA, Walter JM, Joshi N, Anekalla KR, McQuattie-Pimentel AC, Chiu S, Fernandez R, Akbarpour M, Chen CI, Ren Z, Verma R, Abdala-Valencia H, Nam K, Chi M, Han S, Gonzalez-Gonzalez FJ, Soberanes S, Watanabe S, Williams KJN, Flozak AS, Nicholson TT, Morgan VK, Winter DR, Hinchcliff M, Hrusch CL, Guzy RD, Bonham CA, Sperling AI, Bag R, Hamanaka RB, Mutlu GM, Yeldandi AV, Marshall SA, Shilatifard A, Amaral LAN, Perlman H, Sznajder JI, Argento AC, Gillespie CT, Dematte J, Jain M, Singer BD, Ridge KM, Lam AP, Bharat A, Bhorade SM, Gottardi CJ, Budinger GRS, Misharin AV. Single-cell transcriptomic analysis of human lung provides insights into the pathobiology of pulmonary fibrosis. American journal of respiratory and critical care medicine 2019;199:1517–1536.

46. Froom Z, Callaghan NI, Davenport Huyer L. Cellular crosstalk in fibrosis: Insights into macrophage and fibroblast dynamics. The Journal of biological chemistry 2025;301:110203.

47. Kolb P, Upagupta C, Vierhout M, Ayaub E, Bellaye PS, Gauldie J, Shimbori C, Inman M, Ask K, Kolb MRJ. The importance of interventional timing in the bleomycin model of pulmonary fibrosis. The European respiratory journal 2020;55.

48. Cui H, Jiang D, Banerjee S, Xie N, Kulkarni T, Liu RM, Duncan SR, Liu G. Monocyte-derived alveolar macrophage apolipoprotein e participates in pulmonary fibrosis resolution. JCI insight 2020;5.

49. Cui H, Banerjee S, Xie N, Hussain M, Jaiswal A, Liu H, Kulkarni T, Antony VB, Liu RM, Colonna M, Liu G. Trem2 promotes lung fibrosis via controlling alveolar macrophage survival and pro-fibrotic activity. Nature communications 2025;16:1761.

50. Lu W, Teoh A, Waters M, Haug G, Shakeel I, Hassan I, Shahzad AM, Callerfelt AL, Piccari L, Sohal SS. Pathology of idiopathic pulmonary fibrosis with particular focus on vascular endothelium and epithelial injury and their therapeutic potential. Pharmacology & therapeutics 2025;265:108757.

51. Moeller A, Ask K, Warburton D, Gauldie J, Kolb M. The bleomycin animal model: A useful tool to investigate treatment options for idiopathic pulmonary fibrosis? The international journal of biochemistry & cell biology 2008;40:362–382.

52. Peng R, Sridhar S, Tyagi G, Phillips JE, Garrido R, Harris P, Burns L, Renteria L, Woods J, Chen L, Allard J, Ravindran P, Bitter H, Liang Z, Hogaboam CM, Kitson C, Budd DC, Fine JS, Bauer CM, Stevenson CS. Bleomycin induces molecular changes directly relevant to idiopathic pulmonary fibrosis: A model for “active” disease. PloS one 2013;8:e59348.

53. Miura Y, Lam M, Bourke JE, Kanazawa S. Bimodal fibrosis in a novel mouse model of bleomycin-induced usual interstitial pneumonia. Life science alliance 2022;5.

54. Wilson MS, Madala SK, Ramalingam TR, Gochuico BR, Rosas IO, Cheever AW, Wynn TA. Bleomycin and il-1beta-mediated pulmonary fibrosis is il-17a dependent. The Journal of experimental medicine 2010;207:535–552.

55. Kloth C, Gruben N, Ochs M, Knudsen L, Lopez-Rodriguez E. Flow cytometric analysis of the leukocyte landscape during bleomycin-induced lung injury and fibrosis in the rat. American journal of physiology Lung cellular and molecular physiology 2019;317:L109–L126.

56. Marchal-Somme J, Uzunhan Y, Marchand-Adam S, Valeyre D, Soumelis V, Crestani B, Soler P. Cutting edge: Nonproliferating mature immune cells form a novel type of organized lymphoid structure in idiopathic pulmonary fibrosis. Journal of immunology 2006;176:5735–5739.

57. Todd NW, Scheraga RG, Galvin JR, Iacono AT, Britt EJ, Luzina IG, Burke AP, Atamas SP. Lymphocyte aggregates persist and accumulate in the lungs of patients with idiopathic pulmonary fibrosis. Journal of inflammation research 2013;6:63–70.

58. Vuga LJ, Tedrow JR, Pandit KV, Tan J, Kass DJ, Xue J, Chandra D, Leader JK, Gibson KF, Kaminski N, Sciurba FC, Duncan SR. C-x-c motif chemokine 13 (cxcl13) is a prognostic biomarker of idiopathic pulmonary fibrosis. American journal of respiratory and critical care medicine 2014;189:966–974.

59. Cocconcelli E, Balestro E, Turato G, Fiorentu G, Bazzan E, Biondini D, Tine M, Bernardinello N, Pezzuto F, Baraldo S, Calabrese F, Rea F, Sanduzzi Zamparelli A, Spagnolo P, Cosio MG, Saetta M. Tertiary lymphoid structures and b-cell infiltration are ipf features with functional consequences. Frontiers in immunology 2024;15:1437767.

60. Khawaja AA, Chong DLW, Sahota J, Mikolasch TA, Pericleous C, Ripoll VM, Booth HL, Khan S, Rodriguez-Justo M, Giles IP, Porter JC. Identification of a novel hif-1alpha-alpha(m)beta(2) integrin-net axis in fibrotic interstitial lung disease. Frontiers in immunology 2020;11:2190.

61. McCubbrey AL, Allison KC, Lee-Sherick AB, Jakubzick CV, Janssen WJ. Promoter specificity and efficacy in conditional and inducible transgenic targeting of lung macrophages. Frontiers in immunology 2017;8:1618.

62. Stackowicz J, Jonsson F, Reber LL. Mouse models and tools for the in vivo study of neutrophils. Frontiers in immunology 2019;10:3130.

63. Wan Q, Zhang X, Zhou D, Xie R, Cai Y, Zhang K, Sun X. Inhaled nano-based therapeutics for pulmonary fibrosis: Recent advances and future prospects. Journal of nanobiotechnology 2023;21:215.

64. Massaro M, Wu S, Baudo G, Liu H, Collum S, Lee H, Stigliano C, Segura-Ibarra V, Karmouty-Quintana H, Blanco E. Lipid nanoparticle-mediated mrna delivery in lung fibrosis. European journal of pharmaceutical sciences : official journal of the European Federation for Pharmaceutical Sciences 2023;183:106370.

65. Xu L, Ishikawa H, Zhou Y, Kobayashi T, Shozu M. Antitumor effect of the selective hypoxia-inducible factor-1 inhibitors echinomycin and px-478 on uterine fibroids. F&S science 2022;3:187–196.

66. Zeng Y, Wei Y, Chen C, Shen J, Qian J, Ma X, Su M, Zhang T, Yu B, Zhu C, Wang S. Ss-hpt nanocarrier delivery of hif-1alpha-shrna reduces pathological lactylation and alleviates inflammation in a sustained hypoxia mouse model. International journal of biological macromolecules 2025;325:146903.

67. Ruigrok MJR, Frijlink HW, Melgert BN, Olinga P, Hinrichs WLJ. Gene therapy strategies for idiopathic pulmonary fibrosis: Recent advances, current challenges, and future directions. Molecular therapy Methods & clinical development 2021;20:483–496.

68. Ashcroft T, Simpson JM, Timbrell V. Simple method of estimating severity of pulmonary fibrosis on a numerical scale. J Clin Pathol 1988;41:467–470.

69. Seger S, Stritt M, Vezzali E, Nayler O, Hess P, Groenen PMA, Stalder AK. A fully automated image analysis method to quantify lung fibrosis in the bleomycin-induced rat model. PLoS One 2018;13:e0193057.

