## supplemental for "Myeloid HIF-1α Sustains Hypoxic Fibrotic Fronts and Drives Pulmonary Fibrosis"

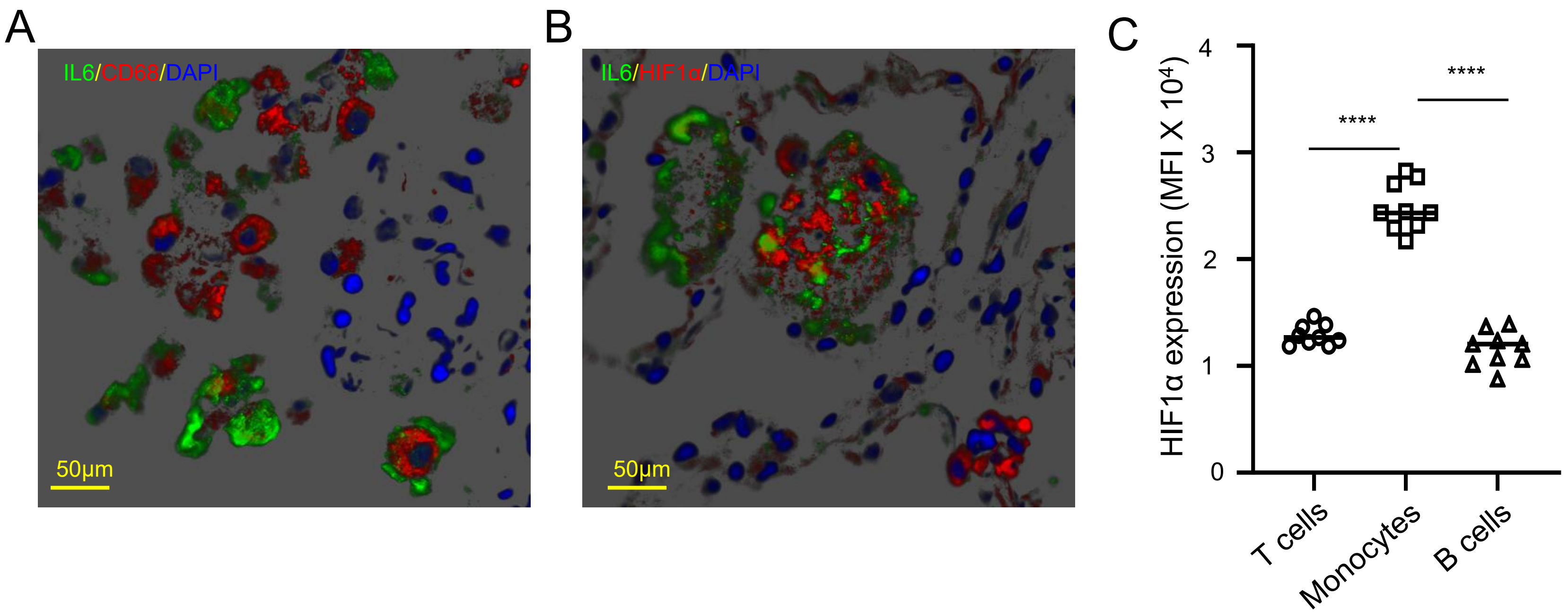


**Fig. S1. Distribution of macrophages, HIF-1α, and IL-6 in human sarcoidosis lung tissue. Related to Figure 1.**
**A.** Representative immunofluorescence image showing IL-6 (green), CD68$^+$ macrophages (red), and DAPI nuclear counterstain (blue). **B.** Adjacent field stained for IL-6 (green), HIF-1α (red), and DAPI (blue). Images are from human sarcoidosis lung specimens. **C.** HIF-1α expression in PBMC subsets from sarcoidosis patients. Intracellular HIF-1α expression was measured by flow cytometry in unstimulated T cells, monocytes, and B cells from PBMCs of 9 patients with sarcoidosis. Expression levels are shown as median fluorescence intensity (MFI). Monocytes exhibited significantly higher HIF-1α expression compared with both T cells and B cells (****$p < 0.0001$, one-way ANOVA with post-hoc test). Data are presented as individual samples with mean ± SEM.

**Table S1.** Demographics and clinical characteristics of the LGRC cohort.

| Characteristic | IPF | Controls |
|---|---|---|
| Number | 123 | 96 |
| Age (years), (mean±SD) | 64.8±8.3 | 63.8±11.2 |
| Males/ Females | 82 (66.7%)/ 41 (33.3%) | 46 (47.9%)/ 50 (52.1%) |
| FVC% predicted | 65.0±16.0 | 95.0±13.0 |
| FEV1%predicted | 71.0±16.0 | 95.0±13.0 |
| DLCO% predicted | 48.0±18.0 | 83.0±17.0 |

Abbreviations: DLCO: Diffusing capacity for carbon monoxide, FEV1: Forced expiratory volume in 1 second, FVC: Forced vital capacity, SD: Standard Deviation.

**Table S2.** The demographic and clinical characteristics of the patient samples

| Patient's Number | Age | Gender | Diagnosis | Grade of disease (GAP STAGE) | GAP SCORE | FVC% | DLCO% | Smoking history |
|---|---|---|---|---|---|---|---|---|
| 1 | 70 | M | IPF | II | 4 | 75 | 66 | 20 PY EX SMOKER |
| 2 | 55 | M | IPF | II | 5 | 36 | 27 | 0 PY |
| 3 | 68 | M | IPF | II | 5 | 48 | 59 | 25 PY EX SMOKER |
| 4 | 64 | M | IPF | II | 4 | 64 | 54 | 15 PY EX SMOKER |
| 5 | 50 | M | IPF | I | 2 | 79 | 47 | 40 PY CURRENT SMOKER |
| 6 | 67 | M | IPF | I | 3 | 80 | 65 | 15 PY EX SMOKER |
| 7 | 69 | M | IPF | III | 7 | 42 | 17 | 45 PY; EX SMOKER |
| 8 | 67 | F | IPF | III | 6 | 42 | 20 | 7.5py; EX SMOKER |
| 9 | 68 | F | IPF | III | 6 | 45 | 28 | NOEN |

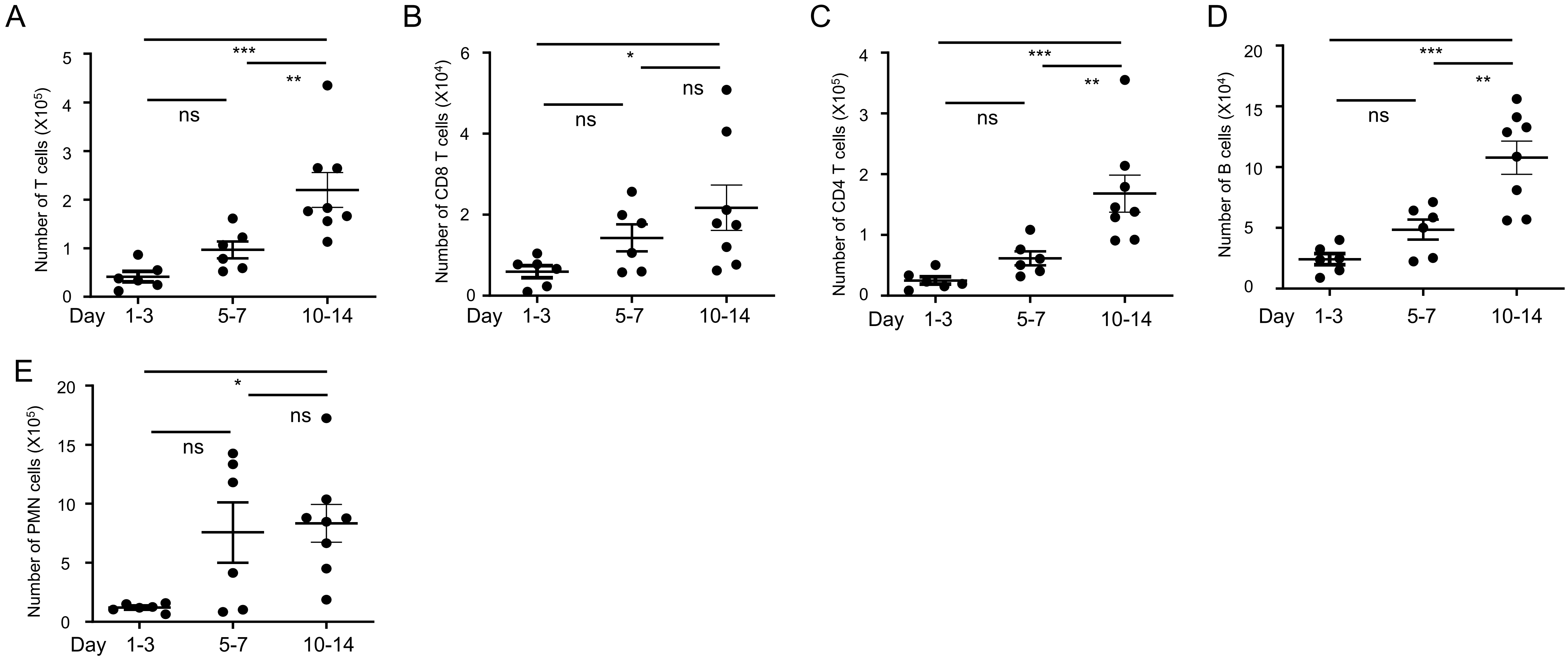


**Figure S2. Temporal dynamics of lymphocytes and neutrophils during bleomycin-induced lung fibrosis. Related to Figure 2.** Numbers of lung **(A)** total $CD3^+$ T cells, **(B)** $CD8^+$ T cells, **(C)** $CD4^+$ T cells, **(D)** $CD19^+$ B cells, and **(E)** polymorphonuclear neutrophils (PMNs) at early (Days 1–3), mid (Days 5–7), and late (Days 10–14) time points after bleomycin injury. Each point represents one mouse; bars indicate mean ± SEM. Statistical comparisons were performed by one-way ANOVA with Tukey's multiple-comparison test. *$p < 0.05$, **$p < 0.01$, ***$p < 0.001$, ns not significant.

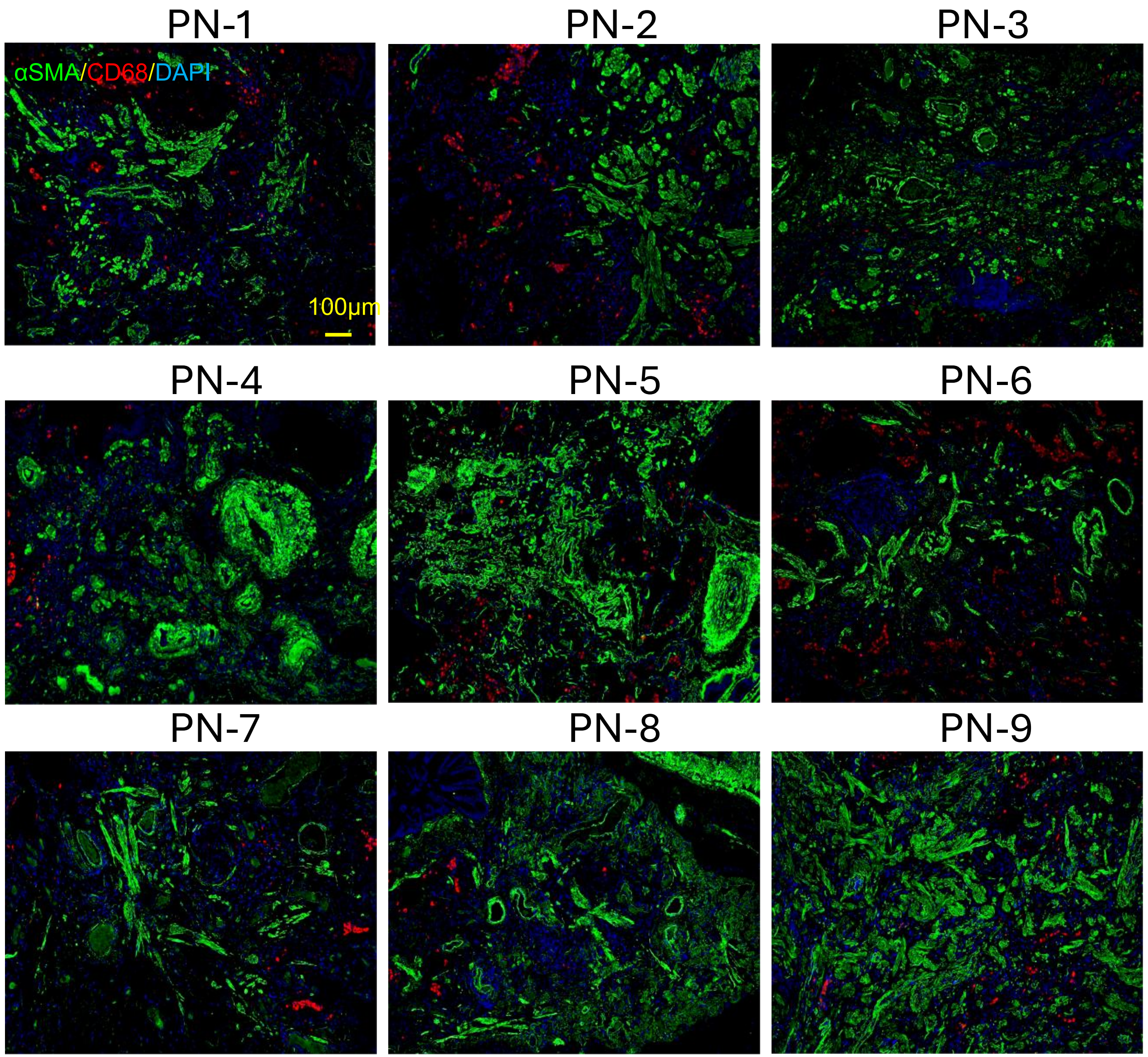


**Fig. S3. Distribution of macrophages and myofibroblasts in additional human IPF lung tissue samples. Related to Figure 3.** Representative immunofluorescence images of αSMA$^{+}$ myofibroblasts (green) and CD68$^{+}$ macrophages (red) with DAPI nuclear counterstaining (blue) from nine human IPF lung specimens (PN-1 to PN-9). Macrophages are predominantly located in peripheral or interstitial regions of fibrotic lesions, with higher abundance in less dense fibrotic areas and reduced presence in mature, collagen-rich scar tissue. Scale bar: 100 μm.

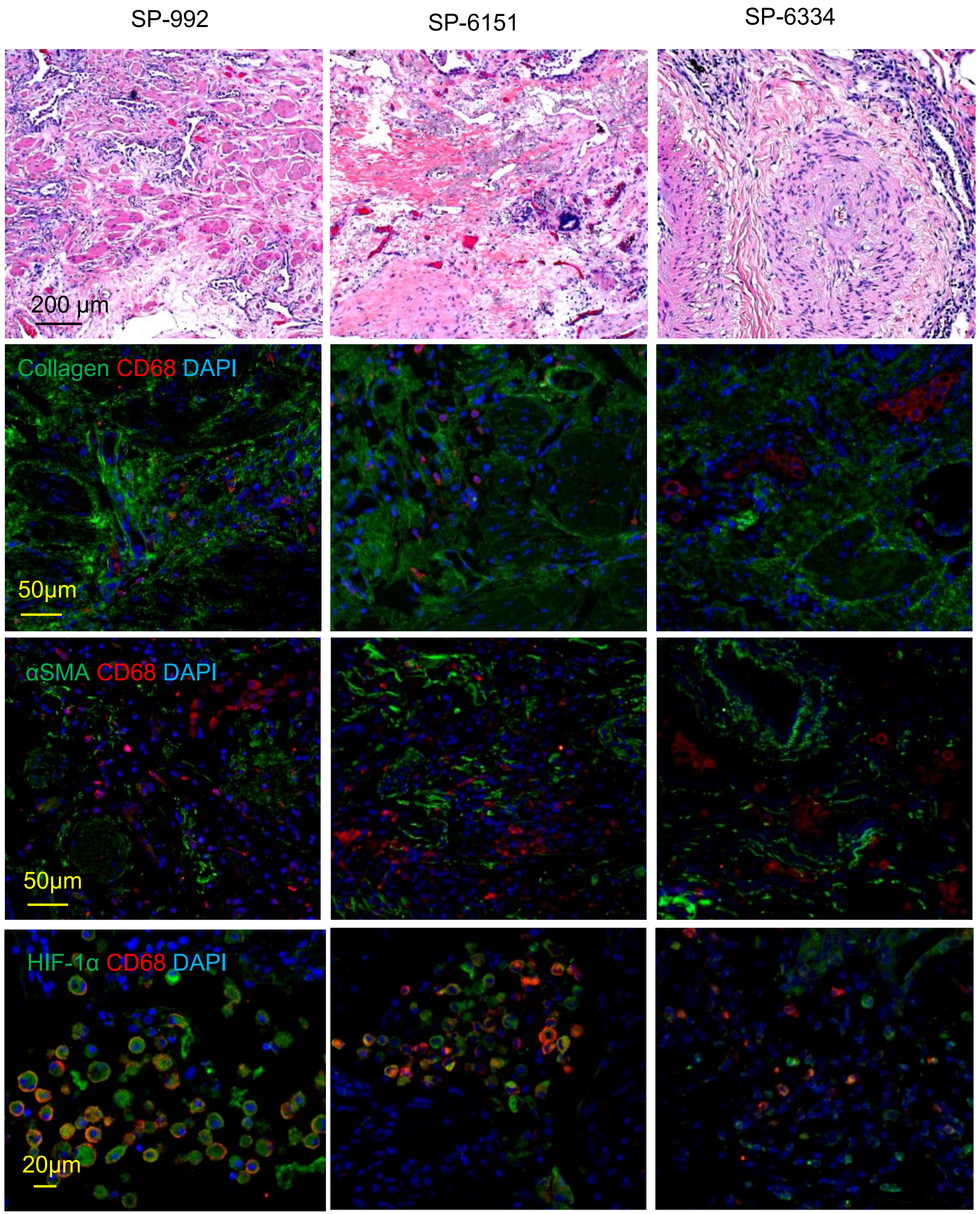


**Fig. S4. Fibrotic remodeling and macrophage-associated HIF-1α in human sarcoidosis lung. Related to Figure 3.** Representative images from three sarcoidosis patients (SP-992, SP-6151, SP-6334). Top panel: H&E staining showing granulomatous/fibrotic pathology. Second panel: Immunofluorescence staining for collagen (green), $CD68^{+}$ macrophages (red), and DAPI (blue). Third panel: Immunofluorescence staining for αSMA (green), CD68 (red), and DAPI (blue) highlighting myofibroblast-rich fibrotic regions and associated macrophage infiltration. Bottom panel: Immunofluorescence staining for HIF-1α (green), CD68 (red), and DAPI (blue), demonstrating HIF-1α accumulation within $CD68^{+}$ macrophages in diseased regions.

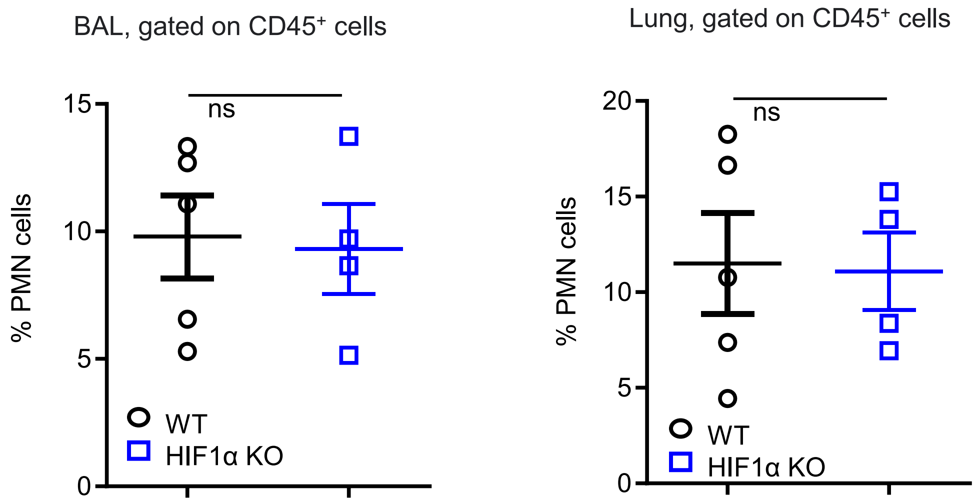


**Fig.S5. Targeted mutation of the *Hif1a* gene didn't influence the number of neutrophils in BAL and lung in BLM-induced mice. Related to Figure 4.**
**A, B.** Flow cytometry analysis of BAL and lung samples from WT or *Hif1a*$^{-/-}$ neutrophils ($CD45^+CD11c^-Gr1^+CD11b^+$) after BLM administration. The percentage of WT and *Hif1a*$^{-/-}$ neutrophils gated on $CD45^+$ subsets in BAL (A) or lung (B) of 14 days post-BLM-administration, shown as mean ± SEM. Data are representative of 3 independent experiments.

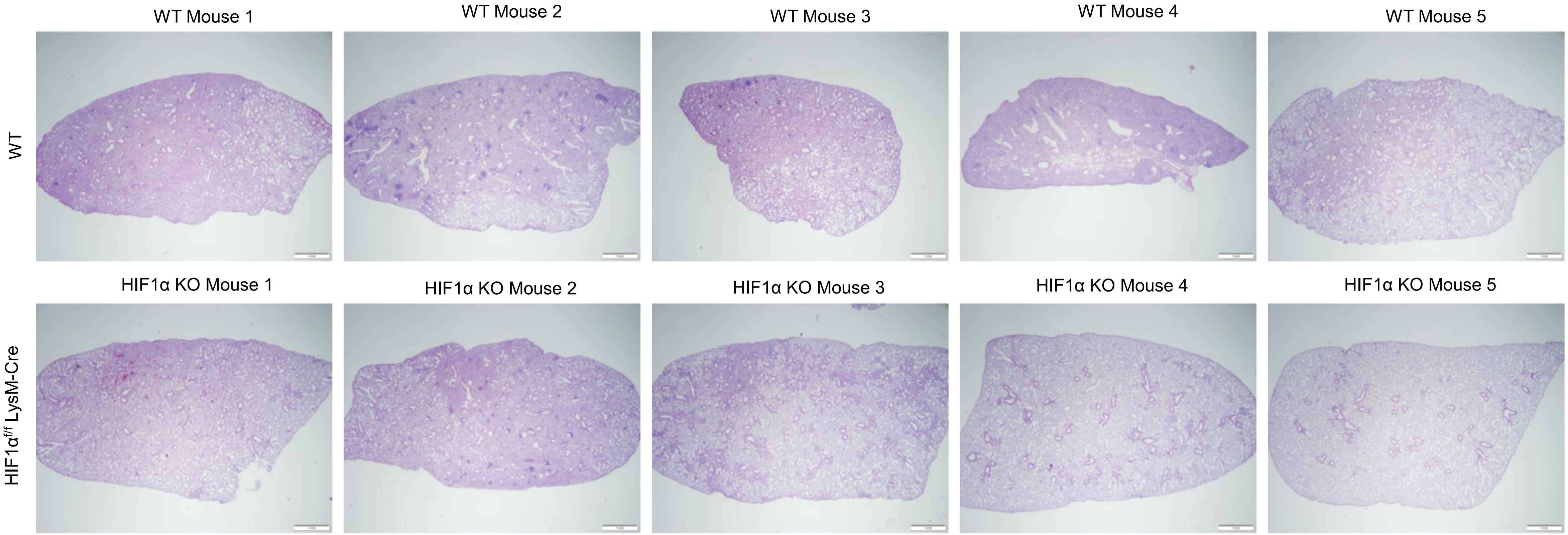


**Fig. S6. Targeted mutation of the *Hif1a* gene mitigates pulmonary fibrosis in BLM-induced mice. Related to Figure 4.** Representative images of H&E-stained lung sections from BLM-induced WT and HIF1α$^{f/f}$ LysM-Cre mice.

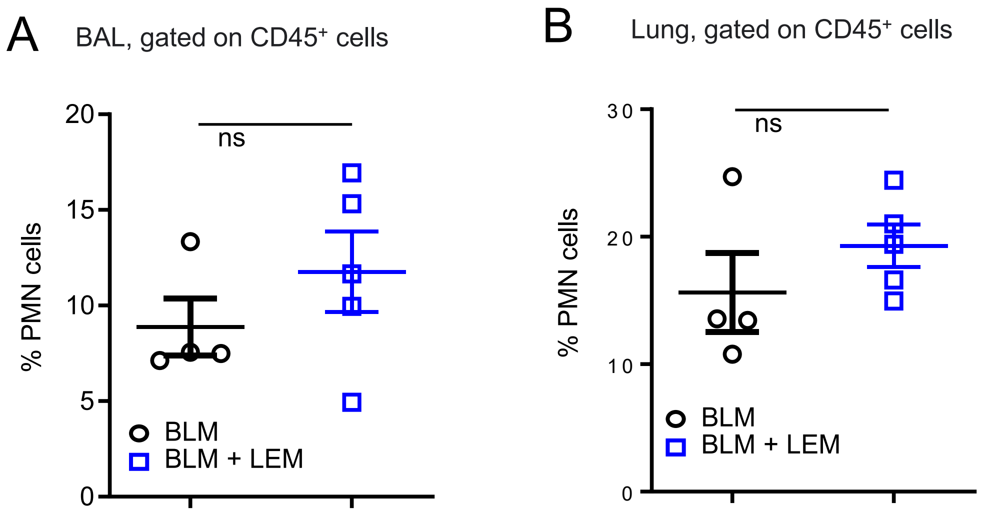


**Fig.S7. LEM doesn't affect the number of neutrophils in BAL and lung in BLM-induced mice. Related to Figure 5.**

**A, B.** Intranasal administrated LEM didn't affect the number of neutrophils in BLM-induced mice. Mice were intratracheally administrated bleomycin on day 0 and on day 3 intranasally treated with 0.05mg/kg of LEM for a cycle of 5 contiuning day with 2-day interval for 2nd cycle. On day 14, BAL and lung tissues were collected. The frequency of neutrophils in BAL(A) and lung (B) gated on $CD45^+$ subsets from LEM- or vehicle-treated mice analyzed by Flow cytometry analysis, shown as mean ± SEM. Data are representative of 3 independent experiments.

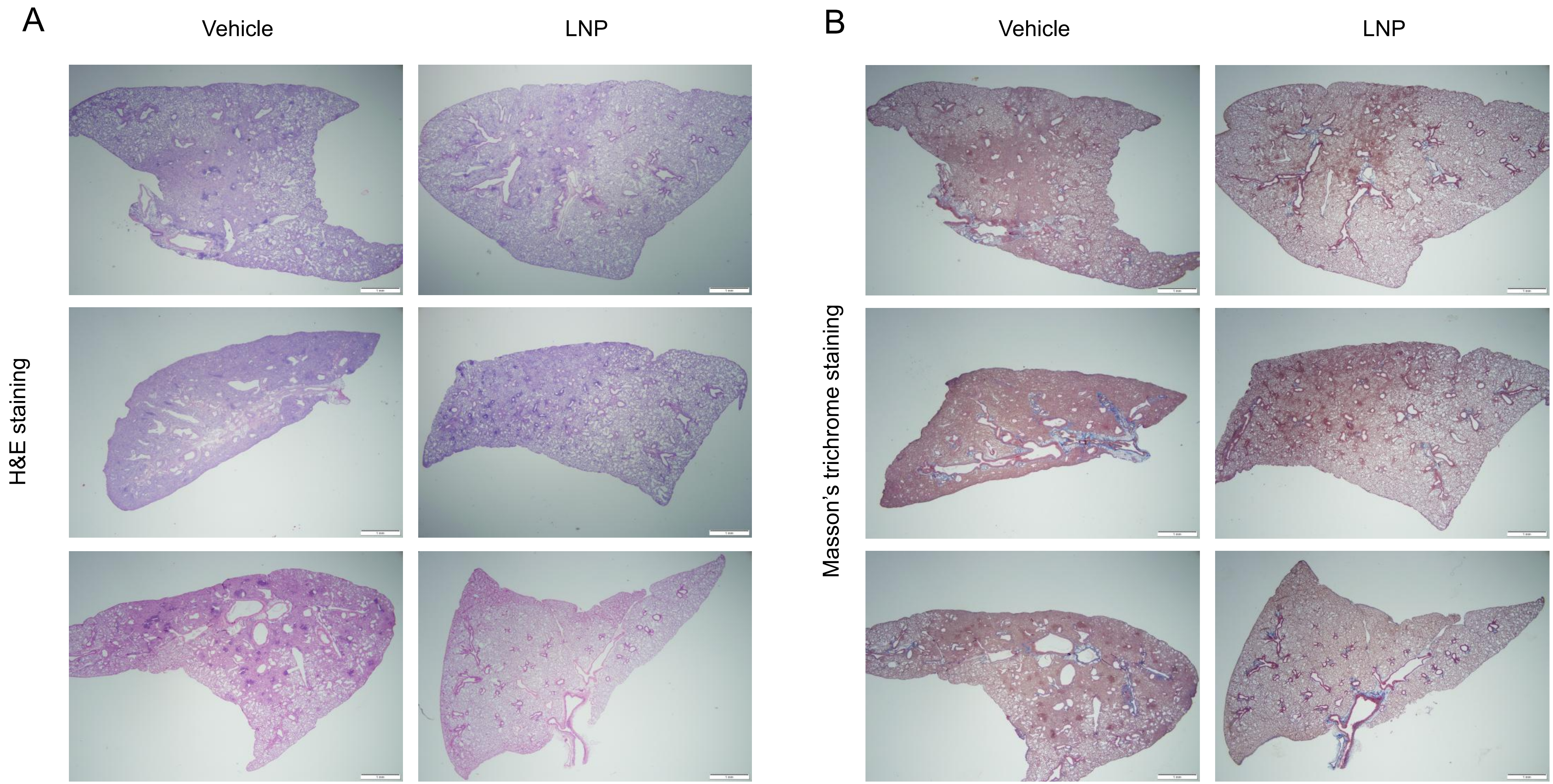


**Fig. S8. Targeting HIF1α with shHIF1α-LNP suppresses fibrosis in a bleomycin-induced pulmonary fibrosis model. Related to Figure 6. A.** Representative images of H&E-stained lung sections from vehicle-treated and shHIF1α-LNP-treated mice. **B.** Representative images of Masson's Trichrome-stained lung sections from vehicle-treated and shHIF1α-LNP-treated mice, highlighting collagen deposition.
